# PLANT MICROTECHNIQUE WITH RESIN - TOWARDS PLANT HISTOLOMICS

**DOI:** 10.1101/2025.11.18.689160

**Authors:** Ivan T. Cerritos-Castro, Araceli Patrón-Soberano, Ana P. Barba de la Rosa

## Abstract

Plant microtechnique is a sequence of skill-intensive histological and microscopy procedures that often yield limited quantitative information. However, it provides the cellular context needed to uncover biomolecular functions. In this work, we developed an easier microtechnique and a novel histolomic approach for the quantitative analysis of histological features. We replaced paraffin with resin as the embedding medium, developed an adhesive treatment for glass slides, and developed a trichrome staining. These improvements provided superior tissue stability and greatly facilitated the skill-dependent steps. Unlike current stainings, our trichrome staining produced a broader color palette and sharply contrasted numerous organelles and ultrastructures in light microscopy. We leveraged these microtechnique advances through image segmentation and quantitative analysis in MATLAB and Adobe Photoshop to measure a wide range of morphometric and compositional features, thereby generating the histolome. To validate this workflow, we applied it comprehensively and systematically to several model plants and calculated their C_4_ Kranz-anatomy level using a combination of characteristic histological features. The histolomes provided new insights into cellular functions and quantitative anatomical differentiation among species. The resin-based microtechnique and histolomic approach will help facilitate, standardize, and make plant histology research quantitative.

**GRAPHICAL ABSTRACT:** 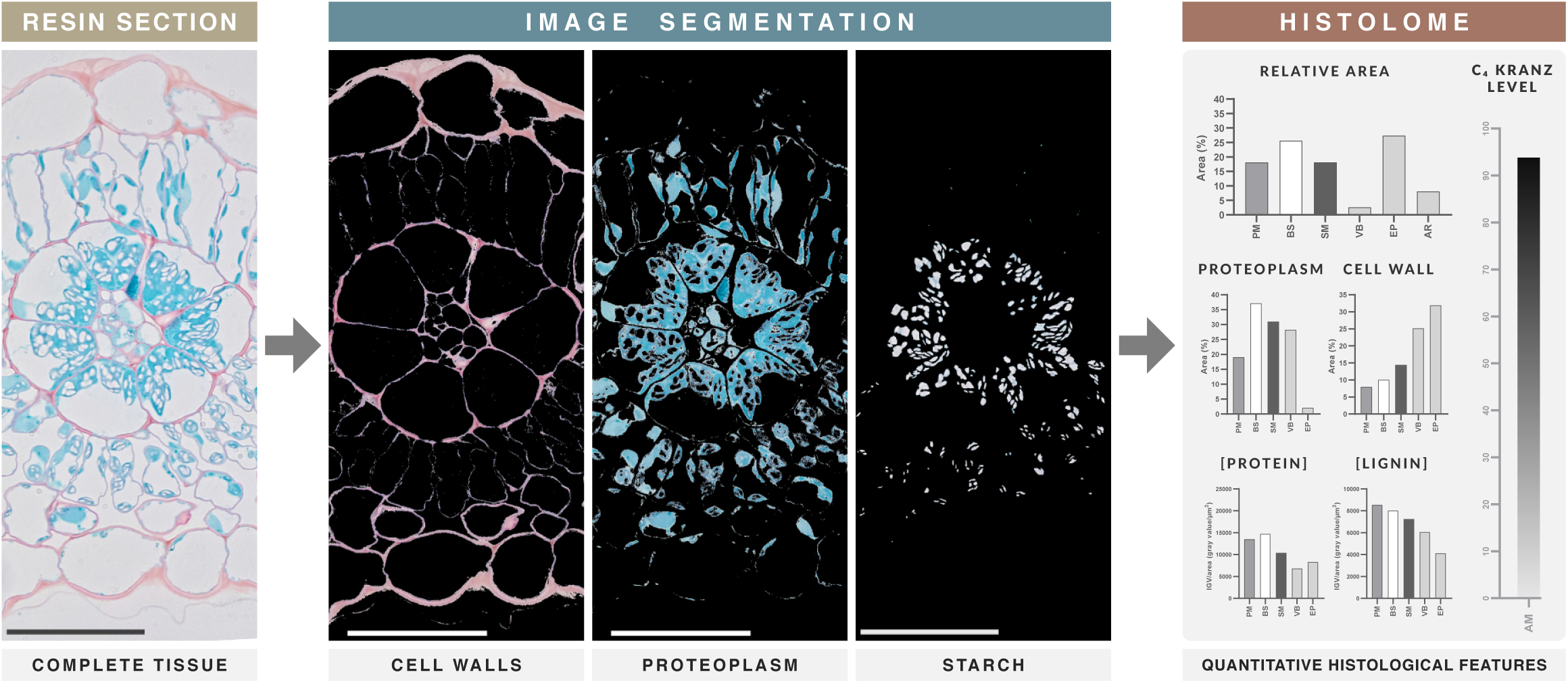

## INTRODUCTION

Tissues arise from the coordinated interaction of millions of ordered molecules. Our understanding of their nature and function has largely relied on molecular disassembly for *in vitro* studies, an approach strengthened by omics techniques that correlate molecular abundances with tissue, organ, and whole-organism phenotypes [1]. However, the location of a molecule is as crucial as its abundance, as spatial distribution is directly related to function and defines tissue composition and morphology [2,3].

Plant histology describes the morphology and composition of tissues and cells [4]. Together with immunostaining, it localizes molecules within their native tissue context [3]. Despite its importance, plant histology remains underutilized in plant research. Two major limitations contribute to this: the high level of skill required for microtechnique procedures and the absence of a harmonized strategy for analyzing micrographs. Compared with *in vitro* molecular methods, plant microtechnique is more artisanal [5]. Classical microtechnique includes fixation, dehydration, paraffin embedding, sectioning, affixing (attachment to glass slides), staining, immunostaining, dehydration, mounting (resin and coverslip placement), and microscopy [6]. Although each step carries the risk of failure, the most skill-dependent steps occur from affixing through mounting. Plant-paraffin sections are fragile: they wrinkle or break during affixing and frequently detach or become scratched during subsequent steps. Moreover, paraffin is removed after affixing, leaving the tissue even more vulnerable to damage.

Replacing paraffin with a more stable, permanent support could facilitate these skill-dependent steps. LR White, an acrylic resin commonly used in transmission electron microscopy (TEM), is also compatible with light microscopy [7]. Its hydrophilic nature allows sections to be analyzed without resin removal, maintaining mechanical support throughout the skill-dependent steps and reducing breakage and scratching. Effective slide cleaning prevents wrinkling, while efficient “gluing” prevents section detachment. Various acidic and oxidizing solutions have been formulated to clean and simultaneously activate slides [8–10]. Activation generates hydroxyl groups that increase wettability. These groups subsequently react with (3-aminopropyl)triethoxysilane (AES) during the “gluing” step, improving slide adhesiveness [11].

Once a section is affixed, staining and immunostaining are the foundational methods of plant microtechnique for assessing morphology, general composition, and protein localization [6]. Johansen’s staining has been the classical method for paraffin sections since 1940 and has undergone several modifications [6,12–15]. It relies on safranin, which stains lignin- and suberin-rich cell walls in magenta, and fast green, which stains cytoplasmic proteins in turquoise [16–18]. Johansen’s staining visualizes the global composition of tissues, whereas immunostaining reveals the spatial abundance of specific proteins [3]. In contrast, resin sections are typically stained only with toluidine blue for light microscopy, which uniformly stains tissues blue [19].

Even after successfully performing microtechnique, many researchers remain uncertain about how to extract quantitative information from micrographs. Histological interpretation is often limited to qualitative descriptions of abnormalities or immunolabel localization. Systematizing histological analysis would allow researchers to fully exploit micrograph information. Tissue features can be described in terms of morphology, morphometry, and composition: morphology refers to qualitative structure; morphometry quantifies number, size, and shape; and composition reflects the overall molecular abundance of cellular structures [4,20,21]. Staining supports boundary-based morphological and morphometric analyses, as well as color-based compositional analyses, which are usually performed manually and visually. However, digital tools—such as color measurement and image segmentation—allow comprehensive numerical characterization that far exceeds what can be achieved by eye [22]. Altogether, an easier and standardized microtechnique combined with a harmonized digital analysis of micrographs would advance plant histology toward a “histolomics” research framework.

Here, we present a resin-based plant microtechnique designed to facilitate the skill-dependent steps, and we introduce a strategy for micrograph analysis: histolomics. We standardized a method for resin embedding, developed a chemical adhesive treatment for glass slides, created a trichrome staining method for resin sections, and integrated it with immunostaining. We then applied this workflow to amaranth and to several C_3_ and C_4_ plant leaves commonly used in plant research. C_4_ plants possess an evolved leaf morphology known as kranz anatomy, which enhances carbon fixation and provides a useful framework for comparative anatomical studies [23]. Finally, we defined and quantified an extensive suite of morphometric and compositional features—the histolome—to systematically characterize and compare plant tissues.

## RESULTS

### Glass slide adhesive treatment

We tested several solutions for cleaning and activating glass slides. Sulphochromic and Piranha solutions produced the best results, yielding the highest wettability and therefore the lowest contact angle. Activated slides were then “glued” with AES. All treatments showed a decrease in wettability and a corresponding increase in contact angle after gluing (Fig. 1a, c). Finally, a resin section was affixed to each slide and subjected to strong washes. Again, sulphochromic- and Piranha-activated slides performed best, retaining the entire section, whereas other treatments resulted in partial or complete detachment (Fig. 1b).

**Figure 1.**
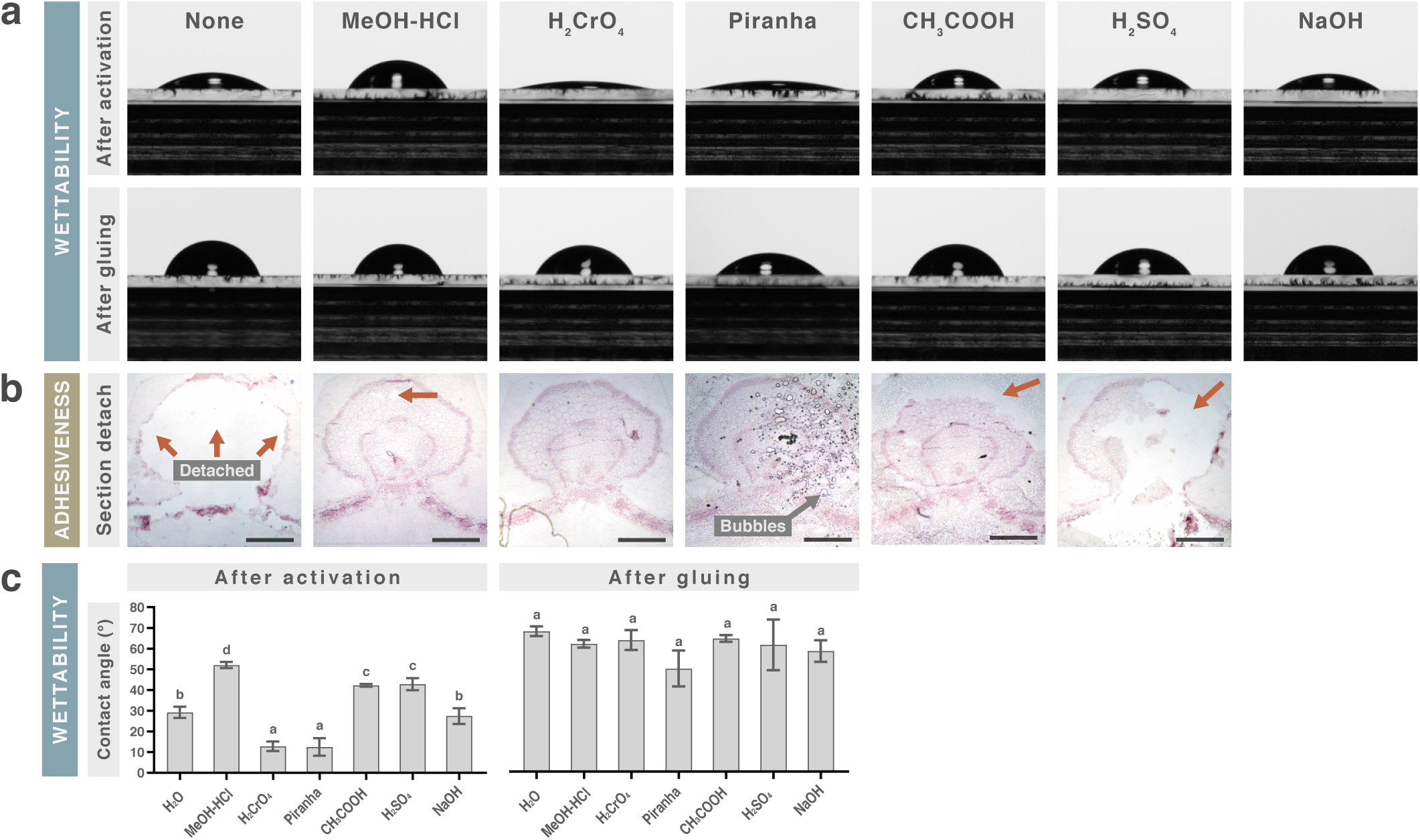
Activation and gluing treatments increase the wettability and adhesiveness of glass slides. a, Contact-angle images after chemical activation and gluing (silanization). b, Resin sections mounted on the different slides after vigorous washing; arrows indicate detachment. Scale bar = 500 µm. c, Mean contact angle (n = 3) ± SD. One-way ANOVA with Holm–Šídák post-tests; different letters indicate p < 0.01. Activation introduces hydroxyl groups that make glass hydrophilic, increasing droplet spreading (wettability) and decreasing the contact angle. These hydroxyl groups subsequently react with AES during the gluing step, improving slide adhesiveness.

### Trichrome staining for resin sections

Classic Johansen’s staining did not work on resin sections; therefore, we reinvented and expanded it into a trichrome method. We tested toluidine blue, Johansen’s, and Coomassie staining and measured the colors acquired by cell walls (CW), protoplasm (PR), and resin background in HSB color space (Hue, Saturation, Brightness). Tissue and cellular structures are shown in Supplementary Fig. 1–2. Hue values depended on the dye, saturation reflected staining performance, and brightness was mainly determined by microscope and camera settings and remained similar across micrographs. Toluidine blue produced monochromatic staining (similar hues), and Johansen’s produced polychromatic but uneven and poorly saturated colors (Fig. 2). Achieving different hues between cell structures is essential for identification and digital segmentation, as is maximizing saturation relative to the unstained resin background.

**Figure 2.**
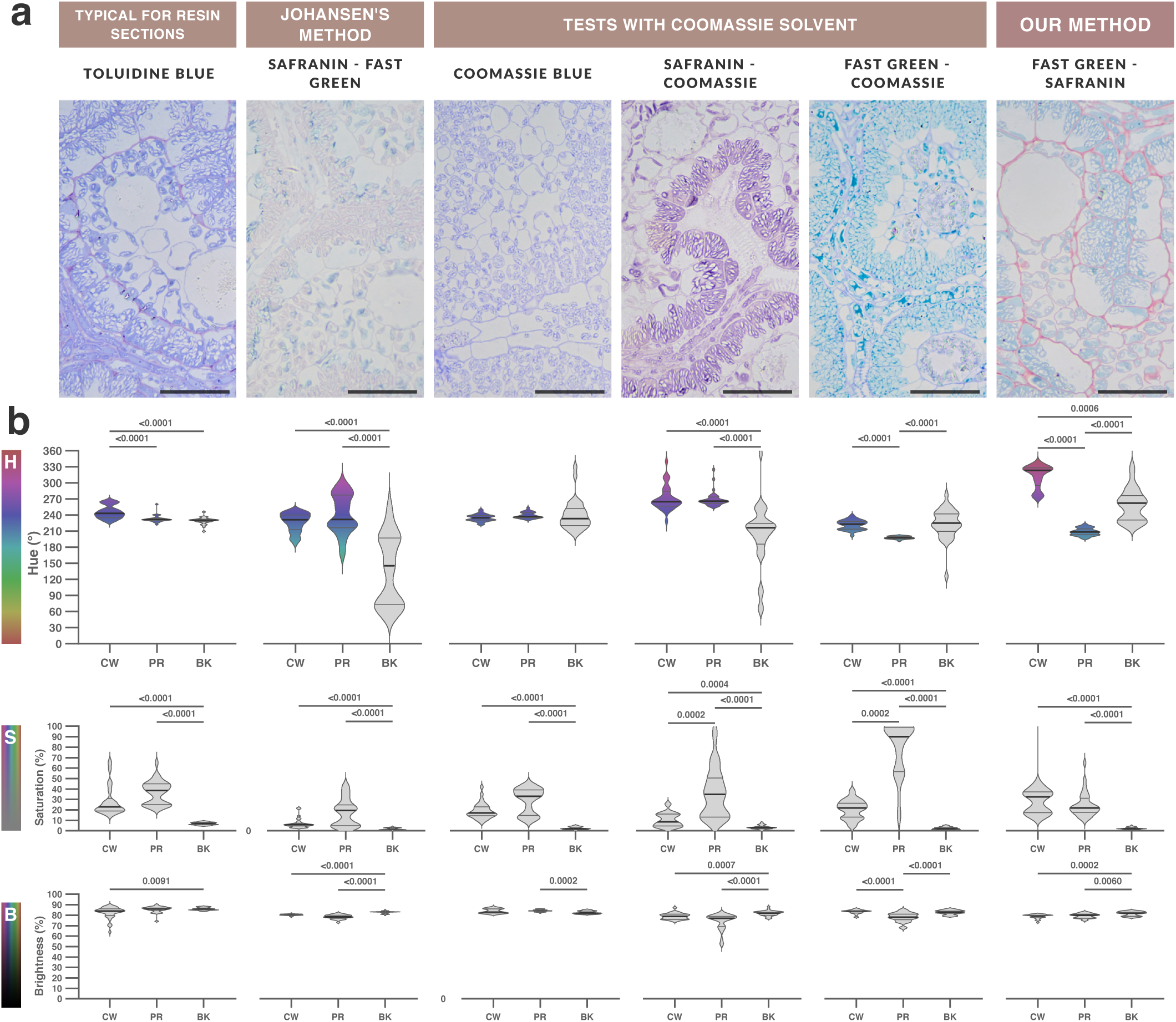
Classic Johansen staining did not work on resin sections. a, Light micrographs of *A. cruentus* paradermal leaf sections stained with different methods. Scale bar = 50 µm. b, Violin plots (n = 30) showing median, interquartile range, and hue ranges (filled when saturation ≤ 15%) for cell wall (CW), protoplasm (PR), and background (BK). Kruskal–Wallis ANOVA with Dunn’s post-test; p < 0.01. HSB graphs correspond to the top micrograph. Hue (H, 0–360°) reflects the spectral position of the dye. Saturation (0–100%) indicates color intensity, and brightness (0–100%) represents luminance. Hue primarily depended on the dye, saturation on staining performance, and brightness on microscope illumination and camera settings.

By testing combinations, we found that Coomassie’s solvent enabled staining of resin sections with Coomassie dye itself as well as with Johansen’s dyes (Safranin and Fast Green), producing polychromatic, uniform, and well-saturated colors (Fig. 2). We therefore reformulated Johansen’s staining and added a third dye—Lugol’s iodine—resulting in a trichrome staining (Fig. 3).

**Figure 3.**
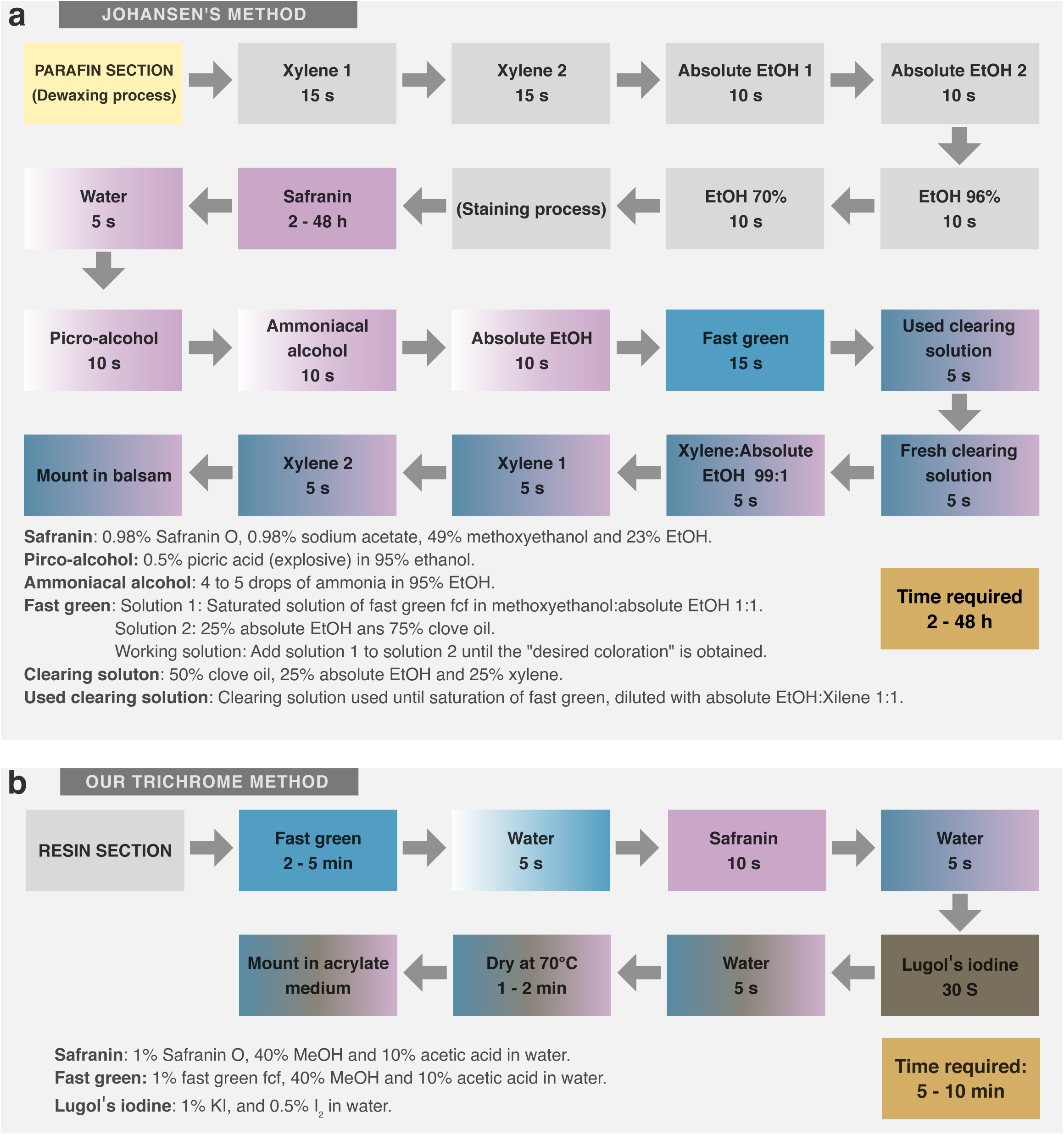
Reinvented Johansen staining for resin sections, converting it into a trichrome method. a, Original 1940 Johansen method (safranin and fast green) for paraffin sections. b, Reformulated trichrome method using fast green, safranin, and Lugol’s iodine for both paraffin and acrylic resin sections. Johansen’s classic method is time-consuming and relies on toxic, explosive, and expensive reagents, and it failed on resin sections. We completely reformulated its dyes and steps, added Lugol’s iodine, reduced processing time and toxicity, and achieved robust staining in both resin and paraffin sections.

Trichrome staining revealed the morphology and general composition of amaranth leaves (Supplementary Fig. 3). Individually, Fast Green stained the protoplasm turquoise, Safranin stained most tissue magenta, and iodine specifically stained starch granules mauve. None of the dyes stained vacuoles or the resin background, and Fast Green and Safranin did not stain starch granules. Because the protoplasm (PR) comprises the cytosol and all organelles, including the vacuole, we introduced the term proteoplasm (PT) to refer specifically to the turquoise (proteinaceous) fraction: the cytosol and all organelles except the vacuole and starch granules.

A synergistic effect emerged in the trichrome combination: Safranin selectively stained cell walls, and both hue and saturation values increased. Proteoplasm shifted from turquoise to blue hues, while cell walls shifted into magenta and reddish hues (Supplementary Fig. 3). We measured cell wall and proteoplasm colors in the epidermis, mesophyll, and bundle sheath. Proteoplasm hues were similar across tissues, but bundle sheath cell walls displayed magenta, reddish, and even orange hues, whereas epidermis and mesophyll cell walls displayed only magenta. The bundle sheath also contained both stained and unstained starch granules, a pattern confirmed by TEM, which revealed electron-dense and electron-lucent granules (Fig. 4).

**Figure 4.**
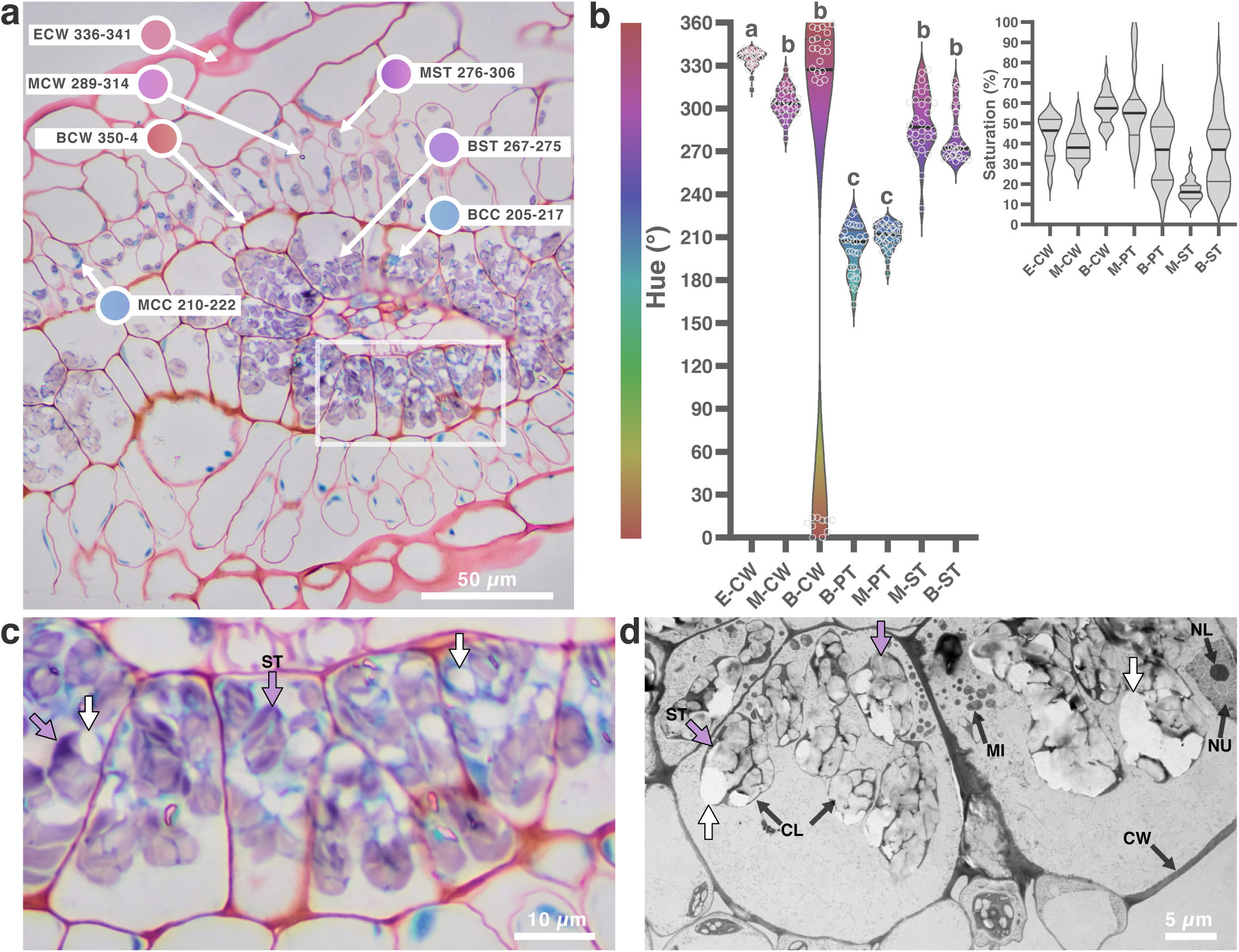
Trichrome staining differentiated cellular composition beyond Johansen dyes. a, Light micrograph of a cross-section of *A. cruentus* leaf stained with the trichrome method; arrows indicate hue ranges of selected structures. b, Violin plots (n = 30) showing median, interquartile range, data distribution, and hue ranges for epidermal CW (E-CW), mesophyll CW (M-CW), bundle sheath CW (B-CW), bundle sheath proteoplasm (B-PT), mesophyll proteoplasm (M-PT), mesophyll starch (M-ST), and bundle sheath starch (B-ST). c, Enlargement of the white box in (a); mauve arrows indicate stained granules and white arrows unstained granules. d, TEM micrograph of a leaf cross-section; arrows correspond to electrodense (mauve) and electrolucent (white) granules. ST, starch; CL, chloroplast; CW, cell wall; MI, mitochondrion; NU, nucleus; NL, nucleolus. Iodine counterstaining shifted the hue of tissues pre-stained with Johansen dyes. The magnitude of the shift was tissue-dependent: M-CW showed minimal change, E-CW shifted toward reddish hues, and B-CW toward reddish to orange, indicating compositional differences in cell walls. Proteoplasm showed negligible hue shift. Some starch granules stained mauve whereas others remained white—paralleling TEM observations and indicating compositional heterogeneity.

The order and timing of dyes were crucial for achieving selective coloration. Fast Green–stained sections could be counterstained with Safranin for 5–10 s; longer exposure caused Safranin to replace Fast Green (Supplementary Fig. 4). Conversely, Safranin-stained sections could not be properly counterstained with Fast Green (Supplementary Fig. 5). Iodine staining—whether used alone or as a counterstain—showed unpredictable intensity, ranging from absent to intermediate or intense, even between serial sections from the same sample on the same slide. We could not identify the underlying cause of this variability. However, within each individual section, staining intensity was always uniform, regardless of whether it was absent, intermediate, or intense (Supplementary Fig. 6).

Some technical considerations are important for optimal staining and micrograph interpretation. The orange hue in bundle sheath cell walls was most evident with a 40× objective and a diaphragm aperture equivalent to 10×. Mounting with Entellan improved sharpness immediately and continued to improve as it dried. Iodine staining may fade over days to months; thus, micrographs should be acquired within one to three days after mounting. Finally, common artifacts associated with resin sections must be recognized for accurate interpretation (Supplementary Fig. 7).

### High-resolution resin-based microtechnique

The resin-based microtechnique allowed light microscopy to resolve small organelles and even ultrastructures typically visible only by TEM. Trichrome staining identified cell walls in magenta to orange, proteoplasm in turquoise, and starch granules in mauve. Differences in proteoplasm saturation enabled identification of cytosol, chloroplasts, mitochondria, nuclei, and nucleoli (Fig. 5; Supplementary Fig. 8–9). An intensely iodine-counterstained paradermal section revealed two layers in the epidermal cell wall—an internal orange layer and an external magenta layer—and additional ultrastructural details such as membranes within vacuoles, chloroplast protein granules, unknown rod-shaped organelles, and vesicles transporting wall material (Supplementary Fig. 10).

**Figure 5.**
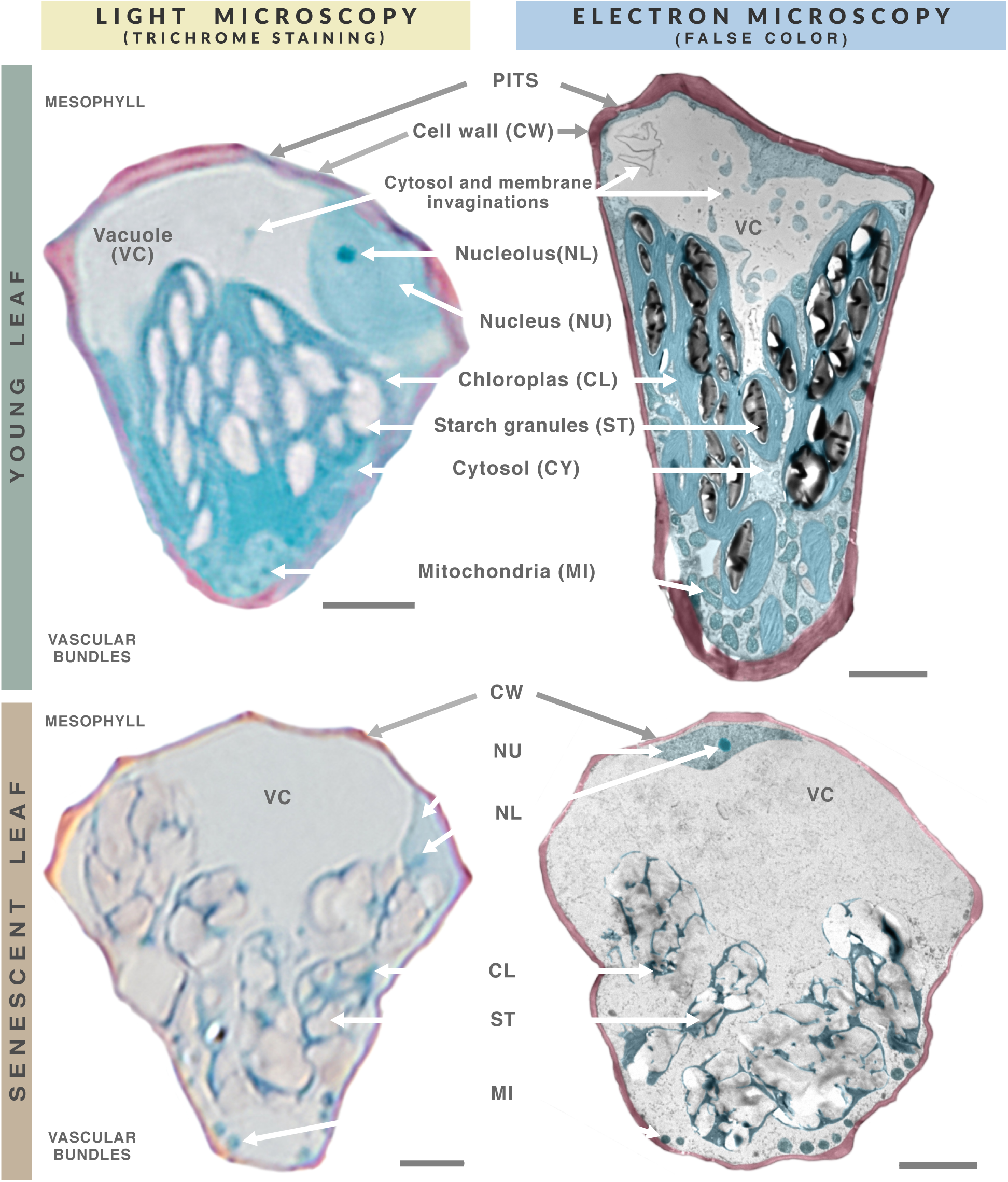
Resin-based microtechnique allowed light microscopy to resolve structures normally requiring TEM. Digitally isolated bundle sheath cells from light and electron micrographs (not the same cell) of young (4-week) and senescent (8-week) *A. cruentus* leaves. Scale bar = 5 µm. High-resolution light micrographs revealed multiple organelles and ultrastructural features validated by TEM. The characteristic PITS-associated wall thinning was visible at bundle sheath–mesophyll junctions. Vacuoles were evident by their lack of color and texture; cytoplasmic invaginations were also detectable. Proteoplasm stained turquoise, and saturation differences allowed discrimination of organelles and cytosol. Although light microscopy did not fully resolve ultrastructures, it enabled reliable identification and tracking across senescence. In senescent cells, cytoplasm was largely lost, chloroplasts lost centripetal arrangement, accumulated more starch, and showed degradation, while nucleus and nucleolus persisted—features detectable both optically and by TEM.

### Histolomic analysis

The resin-based microtechnique required a structured analytical strategy. Toward a histolomics framework, we implemented four steps: visual analysis, color analysis, micrograph segmentation, and data analysis.

First, visual analysis consisted of classical qualitative identification of tissues and morphology (Supplementary Fig. 1–2), with the added identification of organelles and other features enabled by the high resolution of resin sections. Comparing young and senescent amaranth leaves, whole-micrograph differences included decreased proteoplasm, increased starch granules, and rounding of palisade mesophyll cells in senescent leaves. At higher magnification, additional differences were observed: reduced cell wall thickness, appearance of starch granules in palisade mesophyll, and loss of cytosol (Supplementary Fig. 11).

Second, color analysis involved measuring the color of each structure and assigning hue ranges for segmentation (Fig. 4).

Third, micrograph segmentation digitally isolated cellular structures into layers for analysis (Supplementary Video 1). The color contrast provided by the trichrome staining enabled segmentation based on color (composition) and boundaries. We semi-automatically segmented cell walls, proteoplasm, and unstained elements (vacuoles, starch granules, background) based on hue and saturation. Proteoplasm was further segmented into low-saturation elements (cytosol and nucleus) and high-saturation elements (other organelles). Layers clarified tissue composition and spatial patterns, but the thin, discontinuous mesophyll cell walls prevented automated boundary-based morphometric analysis (Supplementary Fig. 12). Therefore, we manually segmented each cell. Segmented micrographs for young and senescent leaves are shown in Supplementary Fig. 13–14.

Fourth, data analysis involved defining and measuring primary and derived morphometric and compositional features. Morphometric features included number, size, perimeter, area, thickness, and circularity; compositional features involved semi-quantitative densitometric analysis of stained molecules (proteins, lignin/suberin, starch, and immunolabels). Senescent leaves showed smaller and more rounded palisade cells, reduced cell wall thickness, decreased proteoplasm, and significantly increased starch content, whereas lignin content did not differ (Supplementary Fig. 15). Intense iodine counterstaining enabled additional segmentation of starch granules, as well as inner and outer wall layers, based on color shift (Supplementary Fig. 16).

### Resin-based microtechnique and histomics validation

The resin-based microtechnique produced comparable results across species and tissues. To test its versatility and the potential of histolomic analysis, we applied it to three C_3_ leaves (*Arabidopsis thaliana, Triticum aestivum, and Glycine max*) and two C_4_ leaves (*Zea mays*, and *Amaranthus cruentus*). The methods worked for all species; dye selectivity matched that observed in amaranth leaves, and iodine staining unpredictability persisted. Histolomic analysis revealed shared fundamental tissues but species-specific differences in morphology, morphometry, and composition (Supplementary Fig. 17–23).

To integrate these data into a physiologically and evolutionarily meaningful metric, we developed a composite feature—the “C_4_ kranz level”—which integrates primary and derived morphometric and compositional features associated with C_4_ Kranz anatomy [23] (Supplementary Fig. 24). *A. thaliana* had the least developed C_4_ Kranz level, and *A. cruentus* had the most developed. We also applied the microtechnique to amaranth seed germination and cotyledon maturation, revealing organelle development and wall thickening (Supplementary Fig. 25–26). Finally, the trichrome staining also worked on paraffin sections, even without deparaffinization (Supplementary Fig. 27).

### Immunostaining of trichrome-stained resin sections

Trichrome staining is compatible with immunostaining, enabling precise protein localization. We performed immunostaining on amaranth leaf sections to detect the RuBisCO large subunit (RbcL). BCIP/NBT development produced an intense blue signal and did not interfere with trichrome staining, resulting in a tetrachrome staining. Micrographs were segmented according to hue ranges, and protein and RbcL concentrations were quantified. RbcL was present in nearly all tissues but was most abundant in the bundle sheath and mesophyll; the bundle sheath contained approximately twice as much RbcL as the mesophyll (Fig. 6).

**Figure 6.**
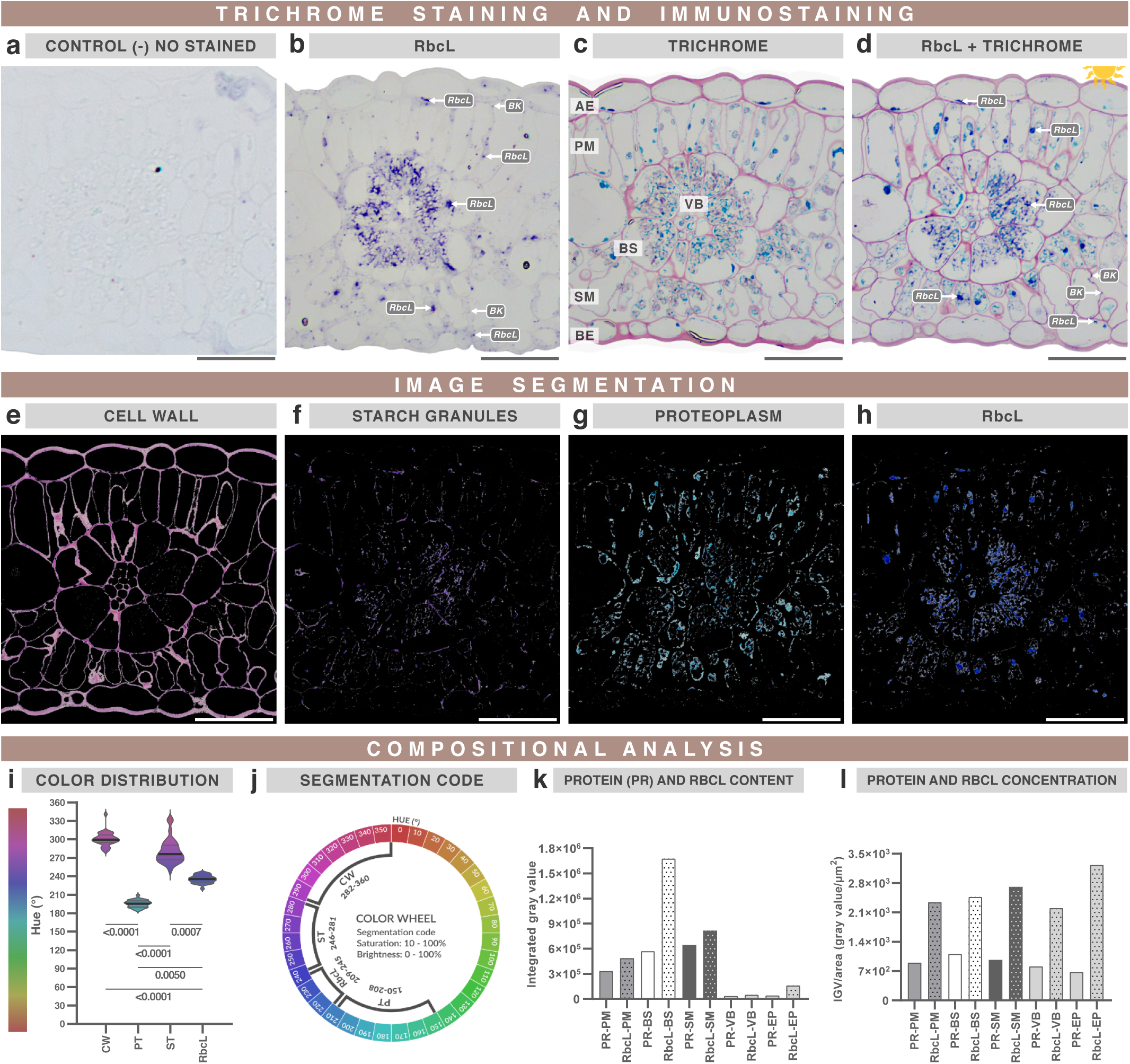
Trichrome staining and immunostaining revealed the precise localization of RbcL in cells and organelles. Light micrographs of *A. cruentus* leaf sections immunostained for RbcL. a, Negative control (no signal). b, Anti-RbcL sample. c, Trichrome staining d, Combined immunostaining + trichrome (“tetrachrome”), AE, adaxial epidermis; PM, palisade mesophyll; VB, vascular bundle; BS, bundle sheath; SM, spongy mesophyll; BE, abaxial epidermis. e–h, Color segmentation of panel (d), isolating the stained tissue components and the RbcL signal. i, Violin plots (n = 30) showing median, interquartile range, and observed color ranges. Kruskal Wallis ANOVA and Dunn’s post-test were performed; differences are shown with p<0.01. j, Segmentation thresholds used in MATLAB. k–l, Densitometric analysis of RbcL and protein content. Immunostaining showed strong RbcL signal and a little background. Immunostaining color did not interfere with trichrome staining colors. Tetrachrome staining improved contrast, facilitating clearer discrimination of RbcL localization compared with immunostaining alone. Although statistical analysis showed significant hue differences among structures (i), starch and cell wall hues partially overlapped (e,f), requiring intermediate segmentation thresholds in MATLAB (j). Densitometry revealed the highest RbcL concentration in bundle sheath cells (k), although relative concentrations were similar across the examined tissues (l).

## DISCUSSION

The resin-based microtechnique simplified the skill-dependent steps of section handling and offered clear advantages over the classic paraffin-based microtechnique. Resin provided continuous mechanical support, prevented breakage and scratching, and preserved tissue structure (Supplementary Fig. 11, 27). Multiple thin and ultrathin sections could be obtained simultaneously for light and electron microscopy or for testing multiple antibodies. Glass-slide activation and AES gluing ensured reliable section adhesion; Piranha- or sulphocromic-activated slides completely eliminated detachment problems (Fig. 1). Because of the toxicity of Cr^6+^, Piranha solution is preferable. Notably, classic methods overlook slide activation and recommend minimal gluing time, which in our experience is insufficient [24]. Activation also cleaned slides deeply, reducing wrinkles and dirt artifacts. However, the main limitations of resin-based microtechnique are the need for an ultramicrotome and the natural wrinkling that occurs during affixing, requiring selection of the best-attached sections.

Our trichrome staining (Fig. 3) reduced steps, time, reagents, toxicity, and variability compared with classical methods [6,12–15]. Johansen’s method involves many steps, large reagent volumes, and toxic or explosive compounds such as picric acid [6]. Our method reduces ≈18 steps to eight, hours to minutes, and nine reagents to six. It also relies on small volumes of accessible, less-toxic reagents, enabling easy disposal and preventing reagent reuse, which improves reproducibility. (Supplementary Video 1).

Trichrome staining revealed the molecular composition of cell structures. Fast Green reportedly stains proteins and is used to quantitatively stain SDS–PAGE gels and nitrocellulose membranes [16,17]. Safranin stains lignin, suberin, nuclei, and chromosomes, although its chromatin specificity is unclear [18,25,26]. Iodine reacts with polysaccharides, with the highest affinity for amylose [27,28]. Our results were consistent with these specificities, except that nuclei, nucleoli, and heterochromatin stained turquoise rather than magenta (Fig. 5). Heterochromatin appeared in *T. aestivum* and *Z. mays*, but magenta staining occurred only in *T. aestivum* (Supplementary Fig. 19–20), indicating that additional molecules may be responsible for safranin affinity. Trichrome staining also distinguished stainable and non-stainable starch granules. Their different colors, densities, and electrolucency indicate two starch types within the same chloroplast, likely differing in amylose content and in their interaction with LR White resin (Fig. 4).

Iodine counterstaining modified the molecular interpretation of the cell wall. It caused a progressive and selective hue shift of safranin-stained walls from magenta to reddish and orange (Fig. 4; Supplementary Fig. 10). Epidermal walls showed two layers—an internal orange layer and an external magenta layer—and segmentation confirmed that all walls retained a small magenta portion (Supplementary Fig. 16). Based on wall composition [29,30], our evidence suggests that orange walls correspond to lignified secondary walls and magenta walls to thinner primary walls. Lignin–iodine interactions can alter absorption spectra [31], explaining the hue shift. However, lignified tracheid rings [4] did not shift (Supplementary Fig. 10). The reactions between iodine and starch granules or wall components raise interesting questions about their compositional differences, and the unpredictability of iodine staining requires further study to standardize.

The resin-based microtechnique resolved tissues, cells, organelles, and ultrastructural details, enabling large-scale histolomic analysis. The smallest structures resolved were membranes inside bundle sheath vacuoles (≈336–560 nm), confirmed as cytosol invaginations by TEM (Fig. 5; Supplementary Fig. 10). These dimensions approach the resolution limit of widefield microscopy (≈200 nm XY; 500–700 nm Z) [32] and overlap with the ultrastructural information typically accessible at low TEM magnification. This “bridge” between light and electron microscopy underpins the histolomic potential of the method: using TEM to understand ultrastructure in detail and then applying high-resolution light microscopy to quantify large areas. Resin sections ≈500 nm thick remains within the depth of field (Z axis) and avoid out-of-focus interference, conceptually similar to confocal microscopy. Thus, the resin-based microtechnique complements high-resolution fluorescent approaches, which visualize only labeled targets.

Histolomics requires standardized analysis and reporting. Micrographs would benefit from an online platform with interactive annotations similar to genomic browsers (Supplementary Fig. 11). Accurate color description is essential; the HSB color space effectively separates dye (hue), staining performance (saturation), and microscope settings (brightness) (Fig. 2). Hue differences allowed segmentation of up to four structures, although color overlap—analogous to fluorescence bleed-through—should be avoided (Fig. 6).

Digital micrographs should be quantified beyond visual inspection. Morphometric features captured differences between C_3_ and C_4_ leaves, though some metrics (e.g., circularity) were insensitive to obvious visual differences (Supplementary Fig. 22). Compositional features captured physiological patterns, such as conserved protein concentration despite reduced proteoplasm during senescence (Supplementary Fig. 15). Primary and derived features can be selected and weighted to study specific phenomena. For the “C_4_ kranz level,” features related to Kranz anatomy [23] were weighted and summed to produce a composite score, enabling comparisons across species and potential correlations with photosynthetic efficiency or C_3_-C_4_ evolution (Supplementary Fig. 24). Similar strategies could quantify tissue responses to stresses, mutations, nutrition, or developmental stages.

More feature descriptors should be developed. Improved shape descriptors are needed, as circularity failed to differentiate tissues. Positional features—such as the location of organelles within bundle sheath cells—could refine functional interpretations. Manual segmentation limits sample size and thus statistical power; deep-learning approaches could automate segmentation and measurement of large areas, fully leveraging the resolution of resin-based microtechnique.

Histolomics preserved the individuality of each cell while enabling large-scale analyses. In *A. thaliana*, we found a previously unreported protein-rich vascular bundle proteinaceous cell (VBPC) with smooth morphology and ≈26% of the vascular bundle protein content (Supplementary Fig. 17), suggesting a specialized function. Across species, tissues, and developmental stages, histolomics revealed both shared and distinctive histological features.

Histolomics also challenged the classification of amaranth as a C_4_ species. RuBisCO usually accumulates mainly in C_4_ bundle sheath cells, where CO₂ is concentrated and thick lignified walls prevent leakage [23]. In amaranth leaves, RbcL, wall thickness, and lignin content increased progressively from palisade mesophyll and spongy mesophyll to bundle sheath. The leaf displayed near-ideal Kranz anatomy, yet its RuBisCO distribution resembled that of C_2_ plants [23].

Whereas omics fields have advanced through rapidly evolving technologies [33], plant histology has relied on almost century-old techniques [6]. Here we reinvented plant microtechnique to simplify methods and produce reproducible micrographs suitable for histolomic analysis. We define histolomics as the large-scale, systematic, high-resolution microscopic study of tissue composition, morphology, and morphometry from ultrastructure to whole tissue. The resulting histolome database of widely studied plant species offers a new perspective for understanding plant structure and its inherent beauty.

## MATERIALS AND METHODS

### Glass slide adhesive treatment

Glass slide adhesive treatment consisted of three steps: washing, chemical activation, and silanization. Slides (2947-25X75, Corning) were first washed by soaking them in 2% soap (Hyclin Plus Neutral, Hycel) for 30 minutes, after which both sides were scrubbed with 15% soap using a streak-free fiber pad (3068257, Scotch-Brite). They were then rinsed with tap water followed by Milli-Q water.

Six activation solutions were tested: MeOH:HCl 1:1 (MeOH–HCl), sulfochromic mixture (K₂Cr₂O₇ ≈ 4% in concentrated H₂SO₄, forming H₂CrO₄), piranha solution (30 ml concentrated H₂O₂ in 70 ml concentrated H₂SO₄), glacial acetic acid (CH₃COOH), concentrated sulfuric acid (H₂SO₄), and 1 M sodium hydroxide (NaOH). The sulfochromic mixture was prepared by forming a paste of K₂Cr₂O₇ with a small amount of water, followed by slow addition of sulfuric acid. All solutions are hazardous; piranha solution and sulfochromic mixture are strong oxidizers that can react violently with organic matter, and the sulfochromic mixture is also highly toxic and must be handled and disposed of according to safety regulations.

Slides were immersed in staining jars containing each solution for 30 minutes at room temperature. A control group received no activation. After treatment, slides were rinsed three times with Milli-Q water, blotted on paper towels, and dried in an oven at 60°C for 30 minutes.

For silanization, dried slides were immersed in 4% (3-aminopropyl)triethoxysilane (AES, 440140, Sigma-Aldrich) in absolute ethanol for 1 hour at room temperature. They were then drained and incubated in 1 mM acetic acid in absolute ethanol for 10 minutes, rinsed with Milli-Q water, and dried again at 60 °C for 1 hour. After the final rinse and drying, slides were stored in dust-free containers to avoid microscopy artifacts.

Water contact angle was measured by placing each slide on a flat, level surface positioned between a backlit flash (Neewer S-400N) and a camera (Canon T5i with EF 100 mm f/2.8L Macro IS USM lens). A 1 µL Milli-Q water droplet was applied to the slide, allowed to stand for 30 seconds, and photographed. Contact angles were quantified in Adobe Photoshop CS6 using the ruler tool.

To evaluate slide adhesiveness under harsh conditions, thin (500 nm) cross-sections of *Amaranthus cruentus* leaf were affixed to the slides at 70 °C for 2 minutes and subjected to three aggressive washing procedures. The first consisted of proteinase K digestion (10 µg/mL in PBS), followed by denaturation at 95 °C for 2 minutes and a 10-second rinse with a wash bottle. The second involved heating in water at 95 °C for 15 minutes followed by a 10-second water-jet rinse. The third involved heating in methanol at 95 °C for 3 minutes followed by a 10-second water-jet rinse. Sections were stained with 0.5% safranin (in water) and imaged. During incubations, slides were covered with microtube lids to prevent drying (Supplementary Video 1).

### Plant material

Seeds of *Amaranthus cruentus* L. cv. Amaranteca (amaranth), *Arabidopsis thaliana* (Arabidopsis), *Triticum aestivum* (wheat), *Glycine max* (soybean), and *Zea mays* (maize) were germinated in 500 g of BM2 HP germination substrate (Berger, Saint-Modeste, Quebec, CA). After 4 weeks, one leaf per plant was harvested. For *A. cruentus*, one leaf was harvested at 4 weeks (young) and one at 8 weeks (senescent). In addition, during germination of *A. cruentus*, seedlings were harvested at different stages—from seed to seedlings with mature cotyledons.

### General sample processing and microscopy

Approximately 3 × 4 mm samples were collected from a primary vein, and the leaf blade. In addition, one seed, one germinating seed, and complete cotyledons at various developmental stages were sampled from *A. cruentus*. Samples were fixed in 3% glutaraldehyde, 2% formaldehyde, and 0.5% Tween 20 in PBS (5–10 times the sample volume) under vacuum for 20 min, then shaken for 1 h, all on ice. Fixation was continued overnight at 4 °C, followed by three PBS rinses, each with shaking for 1 h at room temperature.

Samples were dehydrated in 30%, 50%, 70%, 95%, and absolute ethanol, with shaking for 1 h at each step. Pre-embedding in LR White resin:absolute ethanol (1:1) was performed overnight with shaking, followed by inclusion in pure resin (#14381, Electron Microscopy Sciences, PA, USA) for another night. Pre-embedding can be shortened to 4 h for light, porous samples such as leaves, and maintained overnight for dense samples such as seeds. Typically, all samples were left overnight to ensure complete and homogeneous resin penetration. Samples were then transferred to gelatin capsules #2 (70103, Electron Microscopy Sciences, PA, USA) filled with pure resin and polymerized at 55 °C for 48 h.

For light and electron microscopy, thin and ultrathin sections, respectively, were obtained using a PowerTome PC RMC (Boeckeler Instruments). Samples measuring 3 × 4 mm can be difficult to handle; alternatively, slightly larger samples (≈5 × 6 mm) can be collected, and the excess trimmed with a scalpel or microscissors prior to encapsulation. It is important that the sample does not exceed these dimensions so that it can be properly positioned within the capsule. Additional details on embedding, sectioning, and staining are shown in Supplementary Video 1.

Thin sections were transferred to slides using a Perfect Loop, affixed on a hotplate at 70 °C for 2 min, and subjected to the different procedures described below. Micrographs were acquired using an Axio Imager M2 microscope (Zeiss) and a Canon T5i camera.

Ultrathin sections were placed on Formvar–carbon support film, 150 copper mesh (FCF-150 Cu), and counterstained with 2% uranyl acetate for 10 min, followed by 2% lead citrate for 4 min. Samples were examined with a JEM-200 CX transmission electron microscope (JEOL Ltd.) at 100 kV equipped with a digital camera (SIA, Duluth).

For paraffin embedding, an *A. cruentus* leaf sample was fixed and dehydrated in ethanol as described above, except with two absolute ethanol steps. The sample was infiltrated in absolute ethanol:xylene (1:1) for 1 h, xylene for 1 h, xylene:paraffin (Tissue-Tek VIP 4005 Embedding Medium, Sakura) (1:1) for 1 h at 60 °C, paraffin 1 for 1 h at 60 °C, and finally paraffin 2 for 1 h at 60 °C. Sections of 10 µm thickness (CUT 6062, SLEE, Nieder-Olm, Germany) were obtained, placed in a water bath at 32 °C, collected on slides, and affixed on a hotplate at 60 °C for 2 min. Sections were then stained and micrographs were acquired.

### Trichrome staining for resin sections

Experimental assays. Paradermal sections were stained using different methods. Toluidine Blue O (70103, Ted Pella, CA, USA) was applied for 5 min and rinsed with water. Johansen’s method (Fig. 3a) was performed as described. Coomassie Blue (0.025% Coomassie Brilliant Blue in 40% MeOH and 10% acetic acid) was applied for 5 min and rinsed with water. Safranin (1% safranin O in 40% MeOH and 10% acetic acid) was applied for 10 s, rinsed with water, followed by Coomassie Blue for 10 s and a water rinse. Fast Green (1% fast green FCF in 40% MeOH and 10% acetic acid) was applied for 5 min, rinsed with water, followed by Coomassie Blue for 2 min 30 s and a rinse with water. Finally, Fast Green was applied for 5 min, rinsed with water, then safranin for 5 s and rinsed with water. After the final rinse, slides were dried on a hotplate at 70 °C for 2 min and mounted with acrylic resin (Entellan, Merck) and a coverslip.

Sections shown in Supplementary Figs. 4–6 were stained as indicated in Fig. 3b, with modifications to staining steps, order, and times as described in each figure. Sections in the remaining figures (except Figs. 1 and 2) were stained following Fig. 3b without modification.

### Immunohistochemical staining

Resin sections underwent antigen retrieval with 100 µl of proteinase K at 50 µg/ml in PBS at 37 °C for 30 min. Sections were rinsed with TBST (20 mM Tris, 150 mM NaCl, 1% Tween 20, pH 7.6). Blocking was performed using 200 µl of 1% BSA in TBST (blocking solution) at 37 °C for 30 min. After removing the blocking solution, 200 µl of rabbit anti-RbcL (RuBisCO large subunit) antibody (Agrisera) diluted 1:500 in blocking solution was added. Sections were incubated at 37 °C for 1 h, and then rinsed twice with TBST for 5 min each.

Alkaline phosphatase–conjugated goat anti-rabbit antibody (Merk-Millipore), 200 µl diluted 1:500 in blocking solution, was added and incubated at 37 °C for 30 min. Sections were rinsed twice with TBST, and color development was carried out with freshly prepared BCIP/NBT solution at 37 °C for 2 h. BCIP/NBT solution consisted of 0.02% BCIP and 0.03% NBT in alkaline phosphatase buffer (100 mM Tris, 100 mM NaCl, 5 mM MgCl₂, 0.05% Tween 20, pH 9.5). Finally, sections were rinsed twice with TBST, twice with Milli-Q water, and dried on a hotplate at 70 °C for 2 min.

A negative control was performed without the anti-RbcL antibody, and a positive sample was stained using the trichrome staining. Incubation steps were carried out by covering the section with a microtube lid, as shown in Supplementary Video 1.

### Histolomic analysis - Color measurement

Different areas (n = 30) of the micrographs were measured using the Adobe Photoshop CS6 “eyedropper” tool with a sample size of 3 × 3 pixels, and statistical analyses were performed with GraphPad Prism 9.

### Histolomic analysis - Semi-automatic image segmentation with Matlab R2024a

Micrographs were segmented using the “Color Thresholder” tool in the HSV color space. Segmented images were exported in black and white. To ensure accurate adjustments, HSV values were monitored in the segmentation function and sliders were tuned until the desired values were obtained. Images were exported at a resolution close to the original micrograph (see Supplementary Video 1).

### Histolomic analysis - Manual image segmentation and morphometric and compositional analysis with Adobe Photoshop CS6

Morphometric analysis. A micrograph layer with a layer mask was generated for each element to be measured. The mask density was adjusted to 50%, and each element was manually contoured using the Brush tool. The workspace was calibrated using the “Set Measurement Scale” tool and the micrograph’s scale bar. Selections of each element were retrieved through the layer masks and measured using the “Record Measurements” function. Data were exported and analyzed in GraphPad Prism 9.

Compositional analysis by densitometric quantification of lignin, proteins, and immunolabel. The micrograph was converted to grayscale/negative, and a baseline adjustment was applied. The black-and-white segmented image was resized to match the micrograph and used as a mask. Image analysis was then performed on the masked grayscale/negative micrograph, and the resulting data were saved for statistical analysis in GraphPad Prism 9.

A detailed tutorial on image segmentation and analysis using Matlab and Adobe Photoshop CS6 is presented in the second part of Supplementary Video 1.

## Supplementary material

Supplementary video 1: https://www.youtube.com/watch?v=di2SDqe49rY

## Acknowledgments

Ivan Takeshi Cerritos Castro thanks to CONACYT fellowship 719217, and the grant CNCA.FONCA.04S.04.ACT.PR.ACT.178.18. from the National Fund for Culture and Arts (FONCA), Mexico. The authors thank to Laboratory for Nanoscience and Nanotechnology Research, LINAN, IPICYT.

## Author contributions

**Ivan Takeshi Cerritos-Castro:** Conceptualization, investigation, fund acquisition, methodology, and writing the original draft. **Olga Araceli Patrón-Soberano:** Investigation, methodology, review and editing. **Ana Paulina Barba de la Rosa:** Supervision, funds, resources, review, and editing.

## FIGURES

**SUPPLEMENTARY FIGURE 1.**
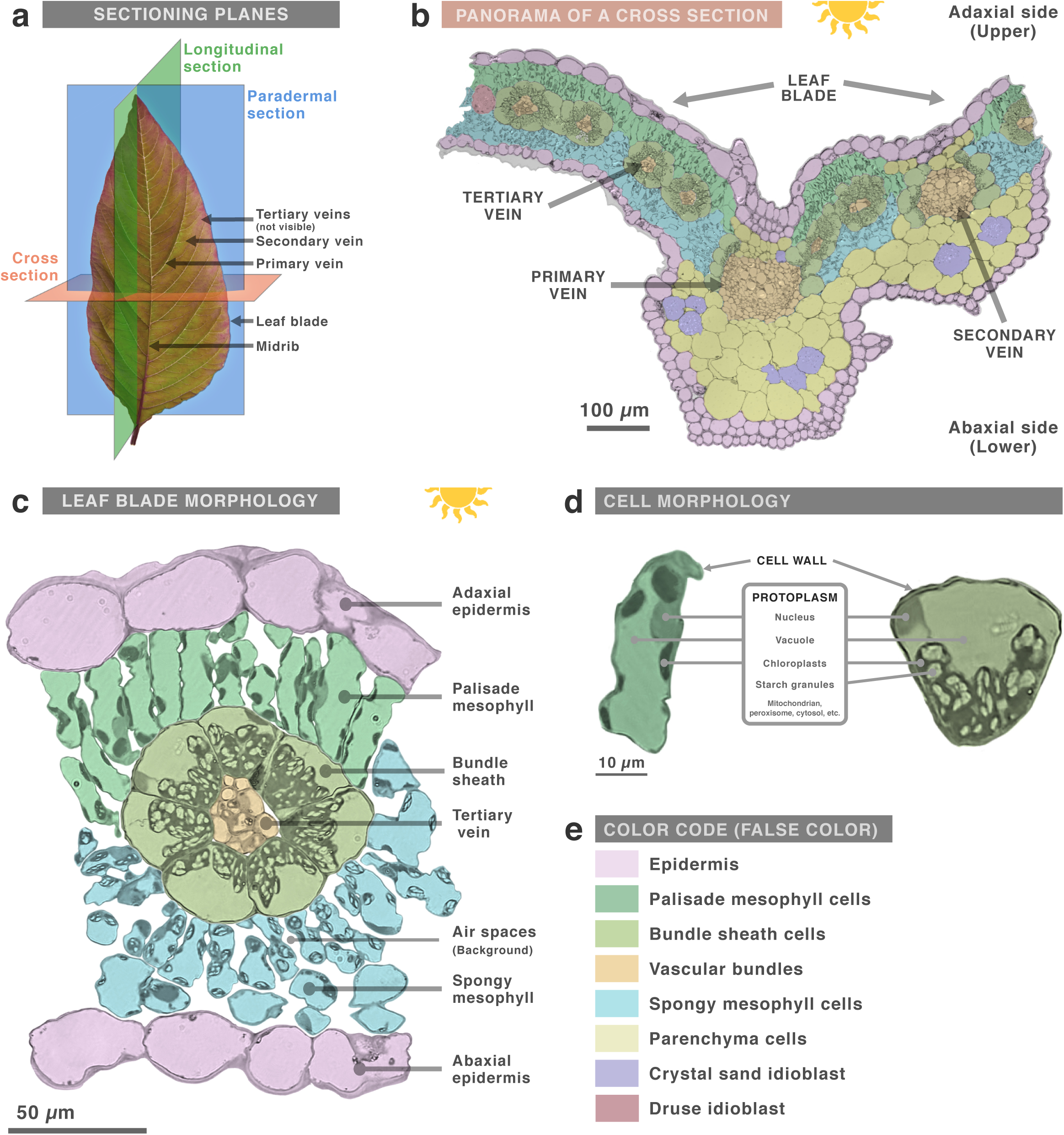
Sectional planes of a leaf and panorama of a cross-section. **a**, Diagram of leaf anatomy and sectioning planes. **b**, Light micrograph of a cross-section of a primary vein of an *Amaranthus cruentus* leaf. The micrograph was colorized and the background digitally removed. **c**, Enlargement of the leaf blade. **d**, Enlargement of palisade mesophyll and bundle sheath cells. **e**, False-color coding to indicate different tissues. Plant leaves consist of a blade supplied by veins that decrease in diameter as they branch away from the midrib. When performed through the midrib or a primary/secondary vein, a cross-section can display all leaf tissues from the adaxial to the abaxial epidermis. Tertiary veins are embedded in the central blade and lack the parenchyma associated with larger vascular bundles; this parenchyma normally produces vein protrusion. Tertiary veins are tightly surrounded by a layer of large bundle sheath cells, which in turn are encircled by a single layer of mesophyll cells, forming a “Kranz anatomy” wreath. Mesophyll is divided into palisade (direct-light exposed) and spongy tissues. Plant cells possess a cell wall, and their protoplasm contains a large vacuole and other organelles. Kranz anatomy and differentiated mesophyll (palisade vs. spongy) are not present in all leaves. Kranz anatomy enhances carbon fixation in C4 plants, whereas plants lacking it are C3.

**SUPPLEMENTARY FIGURE 2.**
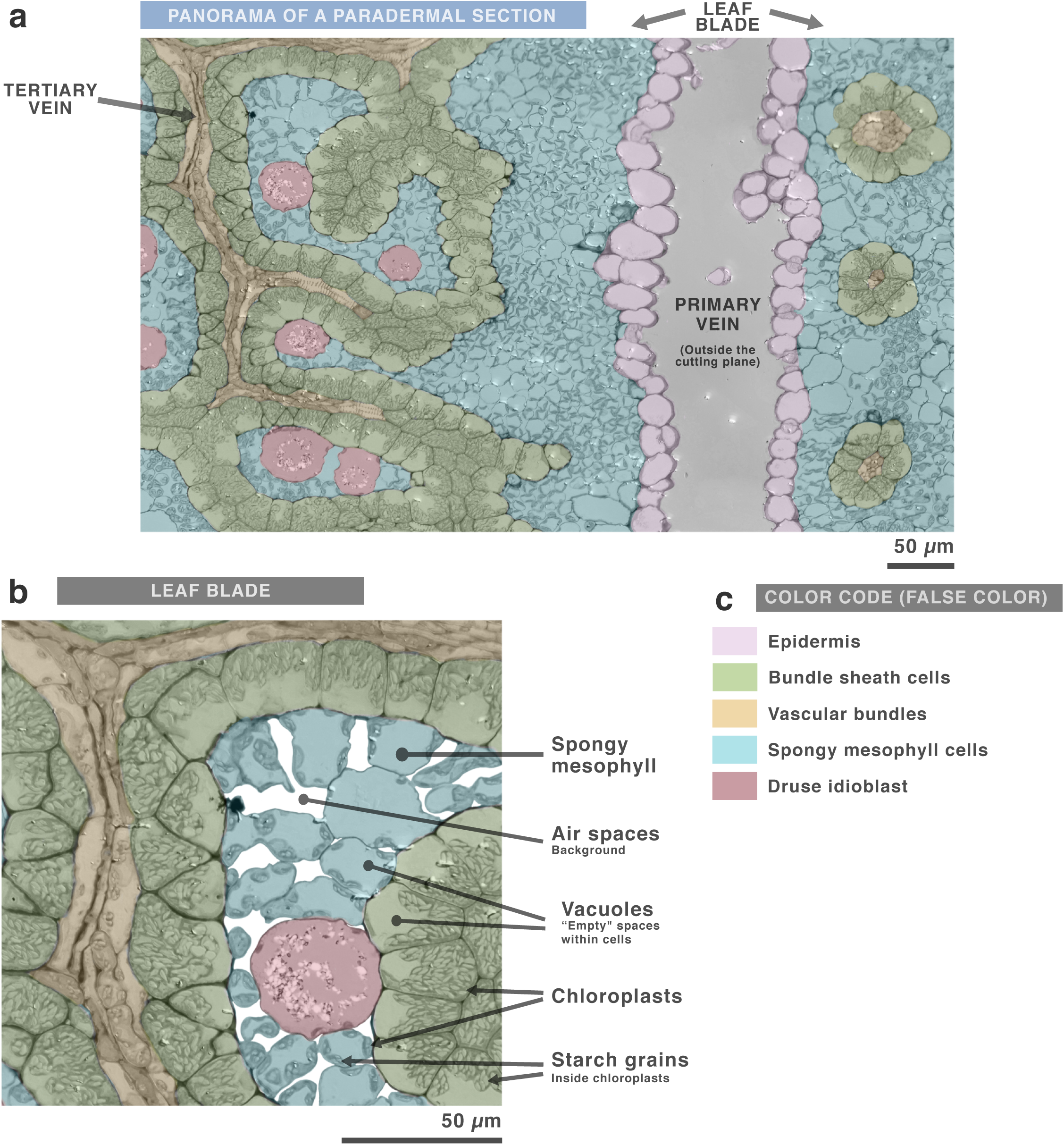
Panorama of a paradermal section. **a**, Light micrograph of a paradermal section of an *Amaranthus cruentus* leaf blade. The micrograph was colorized and the background digitally removed. **b**, Magnification of panel (**a**). **c**, False-color coding indicating different tissues. Paradermal sections are useful for studying the distribution of tertiary veins, minority cell types such as druse idioblasts, and a broader extent of vascular bundles. Cross-sections can show all leaf tissues, but their coverage is limited by leaf thickness. Paradermal sections reveal a wider surface area, though they include fewer tissue types. Because leaves are not perfectly flat, paradermal sections often leave gaps where major veins (midrib, primary, secondary) used to be. Sections taken at mid-thickness are ideal because they contain the greatest cellular diversity. Superficial sections show only epidermal cells, whereas ≈¾-thickness sections capture only mesophyll cells.

**SUPPLEMENTARY FIGURE 3.**
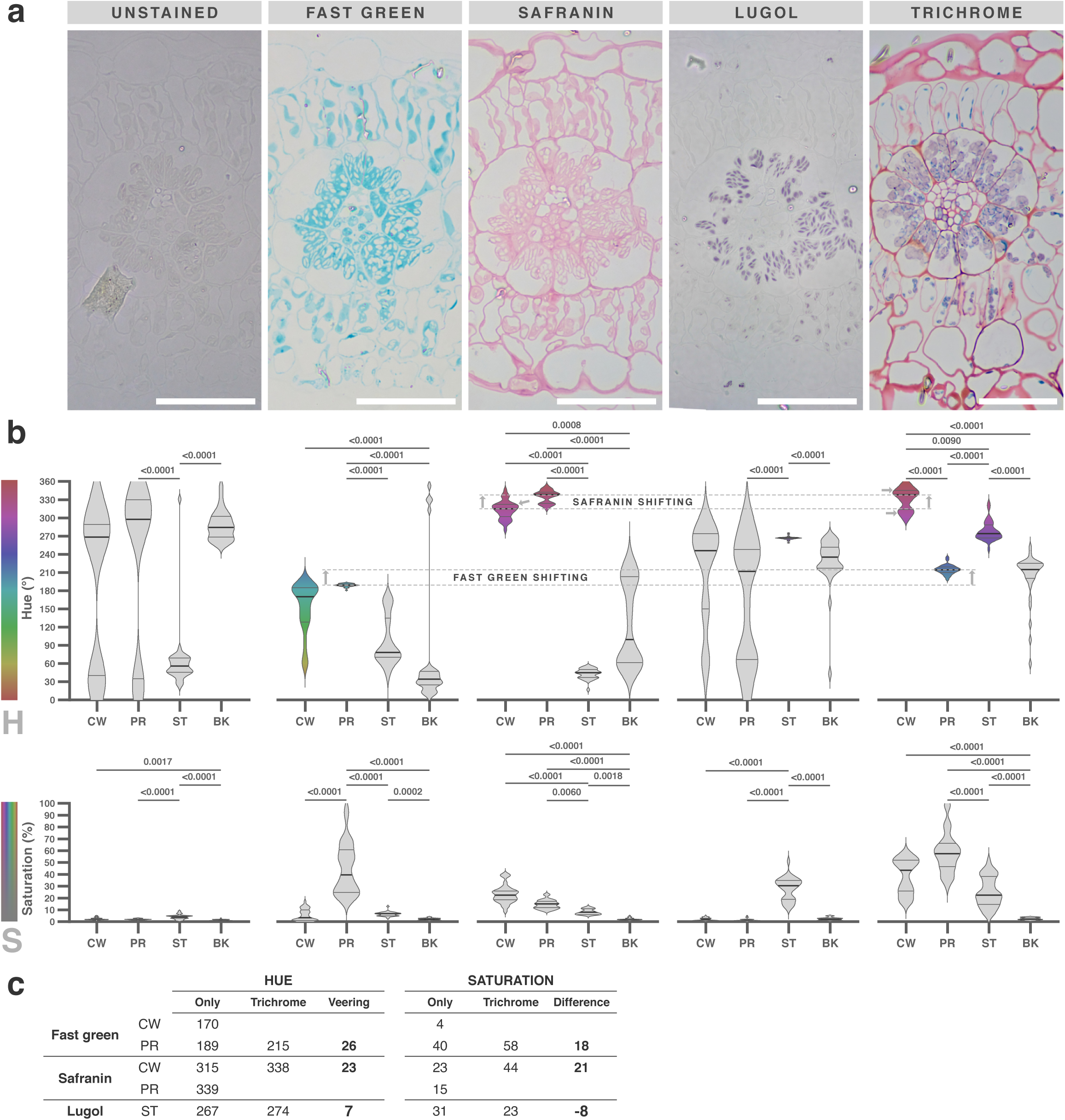
Trichrome dyes show different affinities for cellular structures when used alone or in combination. **a**, Light micrograph of a resin-embedded *A. cruentus* leaf cross-section before staining and after staining with fast green, safranin, Lugol’s iodine, or all three dyes (trichrome). Scale bar = 50 µm. **b**, Violin plots (n = 30) showing median, interquartile range, and color range (filled only when saturation ≤ 15%) for cell wall (CW), protoplasm (PR), starch granules (ST), and background (BK). Kruskal–Wallis ANOVA and Dunn’s post-test were performed; differences are shown at p < 0.01. **c**, Changes in median values for each dye alone compared with the trichrome combination. Resin sections are colorless and visually indistinct prior to staining. Fast green stained the protoplasm turquoise within a narrow hue range (181–193°) but with very broad saturation (19–95%). It also stained the CW across a wide hue range (yellow ≈ 60° to 196°), but with very low saturation (0–17%). Safranin stained both CW and PR in magenta-to-red hues: PR at 315–353° (saturation 9–23%) and CW at 280–346° (saturation 9–40%). Lugol’s iodine stained starch granules mauve within a narrow hue range (259–275°) and a broad saturation range (11–51%). None of the dyes stained the resin background or vacuoles. In the trichrome combination, PR remained turquoise, CW magenta-to-red, and ST mauve. However, slight hue shifts occurred and saturation changed significantly: CW shifted from magenta to red, PR from turquoise to blue, and both increased in saturation. ST showed a broader hue range and reduced saturation.

**SUPPLEMENTARY FIGURE 4.**
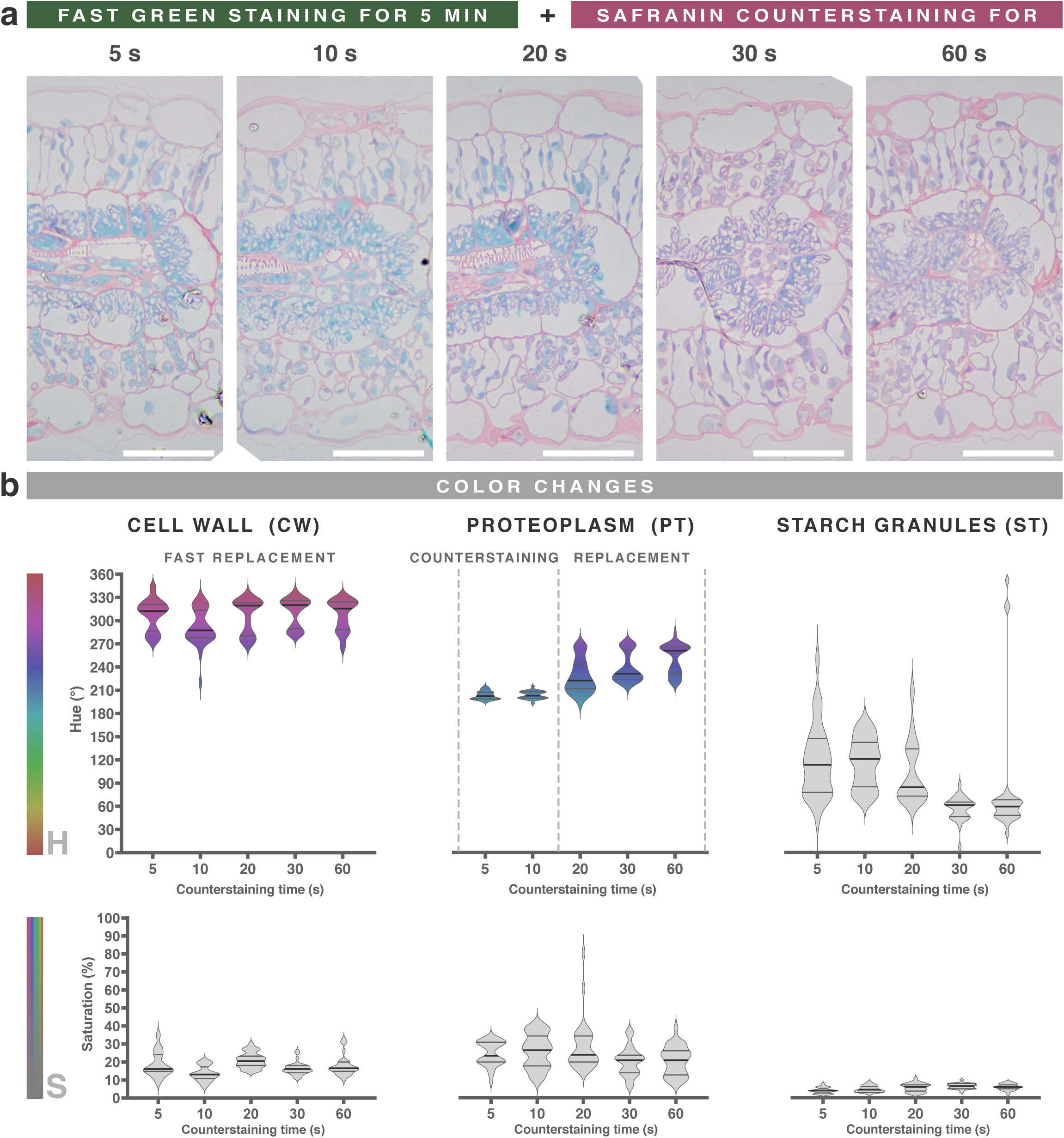
Fast-green–stained sections can be correctly counterstained with safranin for 5–10 s. **a**, Light micrograph of *A. cruentus* leaf sections stained with fast green for 5 min and counterstained with safranin for varying times. Scale bar = 50 µm. **b**, Violin plots (n = 30) showing median, interquartile range, and color ranges for cell wall (CW), proteoplasm (PT), and starch granules (ST) at each counterstaining time. Safranin has strong affinity for lignin and suberin in cell walls, whereas fast green preferentially stains protein-rich proteoplasm. Safranin rapidly and selectively displaced fast green from the CW (counterstaining). After ≈10 s, however, safranin began to displace fast green from its primary affinity region—the proteoplasm—indicating the onset of dye replacement.

**SUPPLEMENTARY FIGURE 5.**
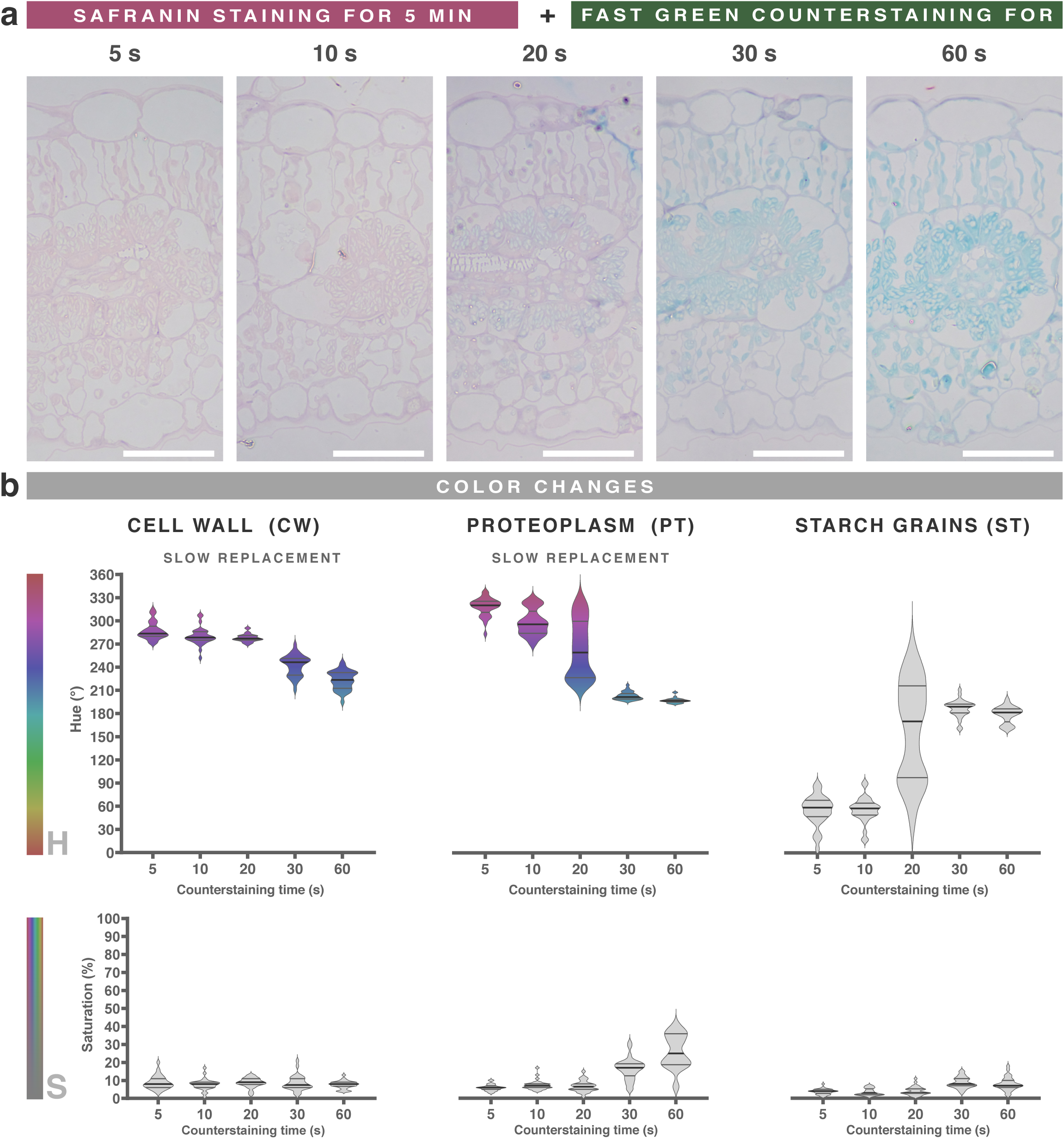
Safranin-stained sections cannot be properly counterstained with Fast Green. **a**, Light micrograph of *A. cruentus* leaf sections stained with safranin for 5 min and counterstained with fast green for varying times. Scale bar = 50 µm. **b**, Violin plots (n = 30) showing median, interquartile range, and color ranges for cell wall (CW), proteoplasm (PT), and starch granules (ST) at each counterstaining time. Safranin exhibits strong affinity for lignified and suberized walls, whereas fast green preferentially binds proteoplasmic proteins. In this order of application, fast green slowly displaced safranin from the proteoplasm—its natural affinity zone—but produced an unsaturated and uneven coloration. After 20 s, fast green also began to remove safranin from the cell wall, generating pale, poorly saturated results. Although this sequence follows Johansen’s classical method, only the reverse order yielded reliable and consistent staining in our resin sections.

**SUPPLEMENTARY FIGURE 6.**
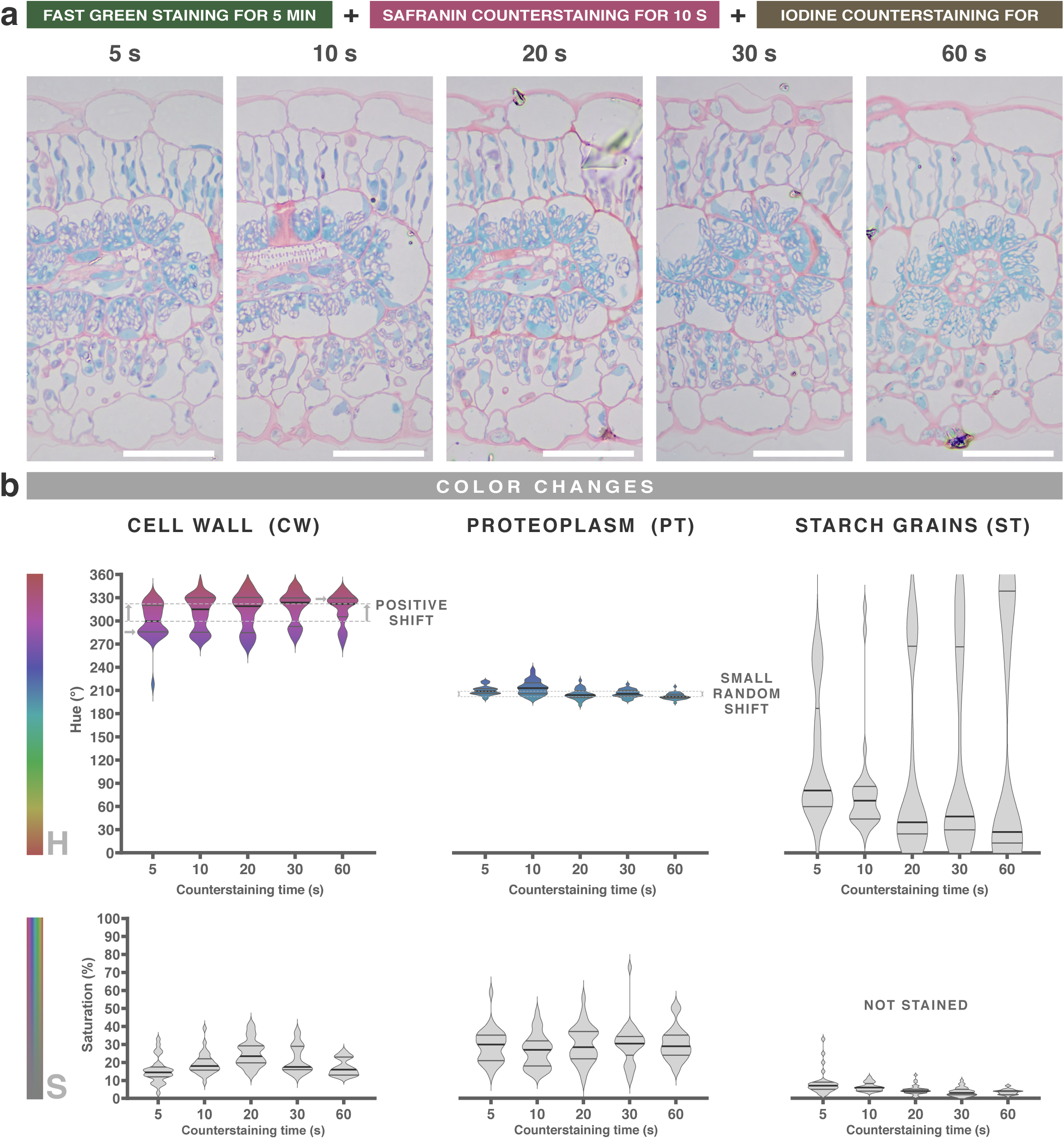
Iodine counterstaining produces hue shifting in stained sections. **a**, ligth micrographs of *A. cruentus* sections stained with fast green (5 min), counterstained with safranin (10 s), and then exposed to iodine for different durations. **b**, Violin plots (n = 30) showing median, interquartile range, and color ranges for cell wall (CW), proteoplasm (PT), and starch granules (ST) at each counterstaining time. Iodine exhibits strong affinity for starch granules and induces hue shifting most noticeably in safranin-stained walls. Iodine staining—either alone or as a counterstain—produced unpredictable intensities in starch granules, ranging from absent to intermediate to intense deposition. This variability occurred independently of time and even among sections mounted on the same slide; however, when staining did occur, it was homogeneous within each section. In the examples shown, starch granules were not stained, but cell walls exhibited clear time-dependent shifting, and the proteoplasm showed mild and random shifts. The underlying cause of inconsistencies in iodine staining remains unknown, requiering numerous replicates to achieve successful results.

**SUPPLEMENTARY FIGURE 7.**
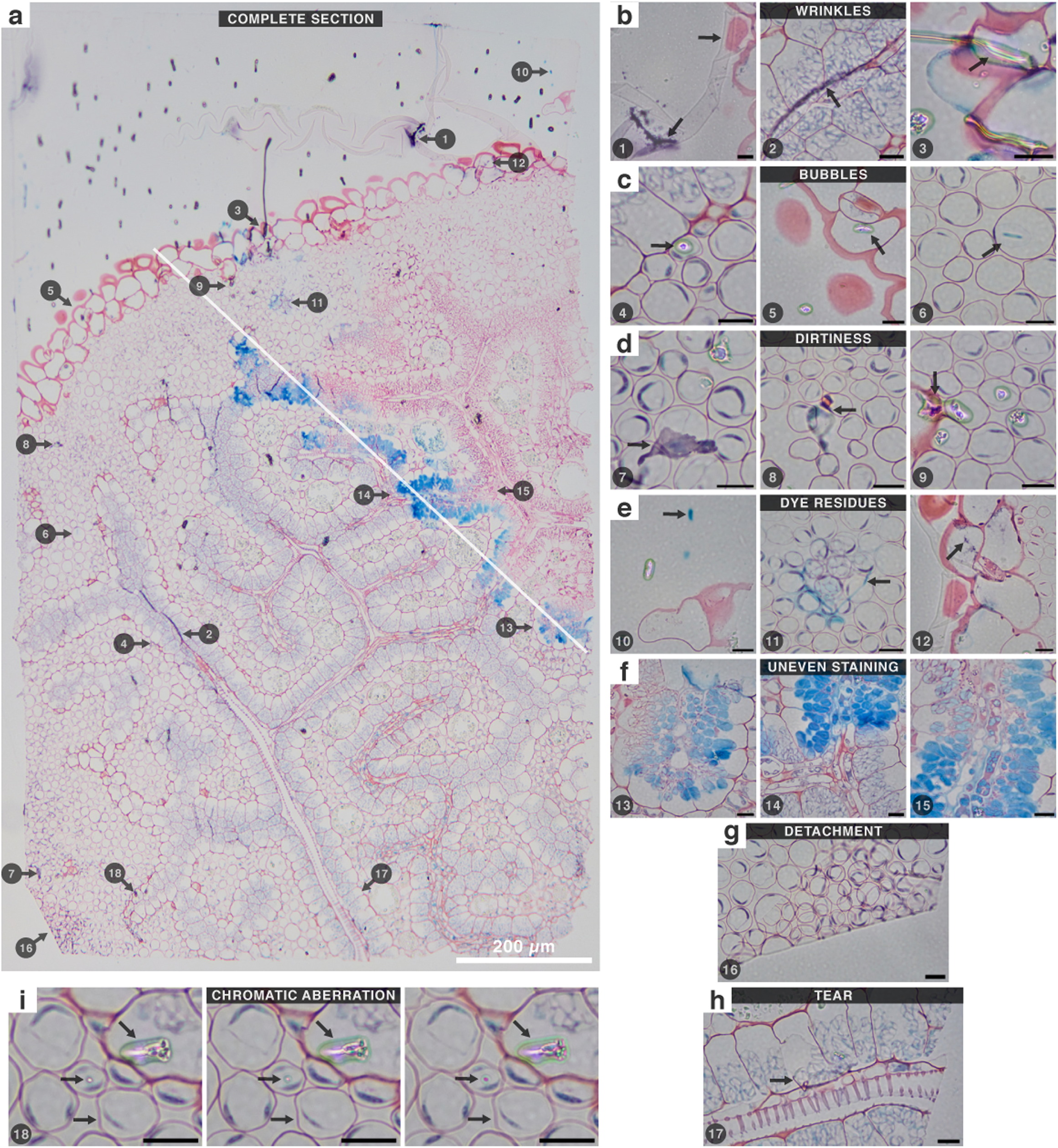
Common artifacts on stained resin sections. **a**, Paradermal section of an *A. cruentus* leaf with magnified insets illustrating recurrent artifacts. (**b**) Wrinkles are the most frequent and difficult issue to avoid, often producing staining irregularities or excessive background in immunostaining. (**c**) Bubbles represent small wrinkles and typically display birefringence, though not always, making them easily mistaken for cellular structures. (**d**) Dust and debris generate localized spots, wrinkles, and bubbles, emphasizing the importance of strict cleanliness when handling specimens. (**e**) Dyes can leave residues when unfiltered, or form precipitates when interacting with other dyes, creating granular deposits. (**f**) Inhomogeneous staining often results from wrinkles, bubbles, or dirt; however, as observed for iodine, it can sometimes occur without any identifiable cause—even in areas perfectly adhered to the slide. (**g**) Conversely, regions partially detached from the glass may stain uniformly. (**h**) Resin sections are also prone to tearing, which distorts tissue morphology. (**i**) Finally, both artifacts and the tissue itself may shift in color depending on the focal plane due to birefringence or chromatic aberration. A consistent focusing criterion is therefore essential for accurate interpretation.

**SUPPLEMENTARY FIGURE 8.**
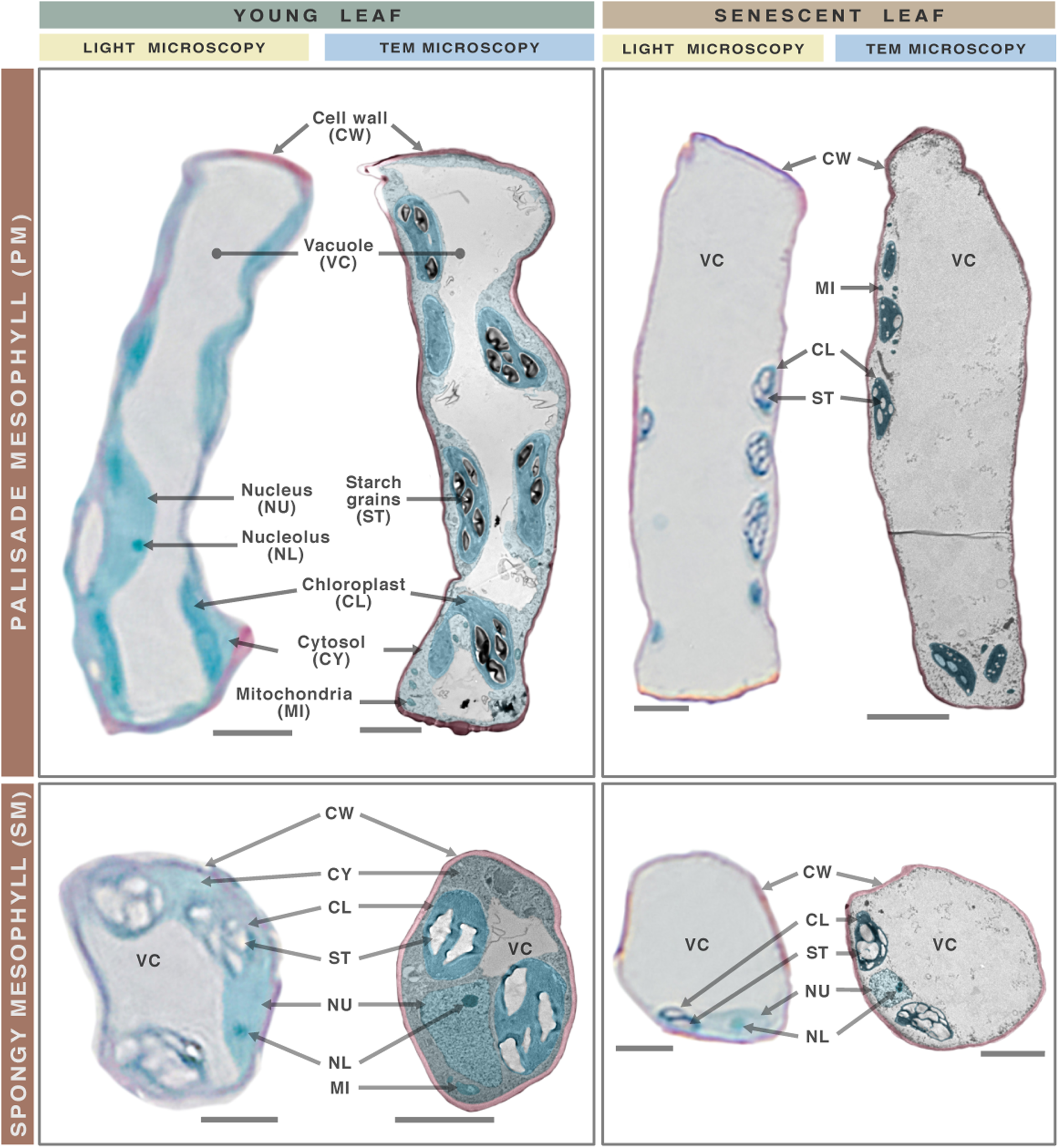
Resin-based microtechnique resolution: mesophyll cells. Light and electron micrographs of digitally isolated palisade and spongy mesophyll cells from young (4 weeks) and senescent (8 weeks) *A. cruentus* leaves are shown (cells are not the same). Light micrographs display trichrome staining, whereas TEM images are false-colored to mimic the same palette. High-resolution resin sections reveal numerous organelles and ultrastructural features confirmed by TEM. The vacuole, identifiable by its lack of color and texture, occupies extensive intracellular volume. The proteoplasm appears turquoise, and variations in saturation reflect different organelles visualized ultrastructurally. Larger or more abundant organelles are more frequently captured because they are more likely to lie within the section plane; smaller or rarer ones—such as the nucleus—often fall outside it. Young leaves show a well-defined cytosol, whereas senescent leaves have lost cytosolic organization and display reduced organelle size. Additionally, chloroplasts in mesophyll cells exhibit variable starch granule accumulation across developmental stages.

**SUPPLEMENTARY FIGURE 9.**
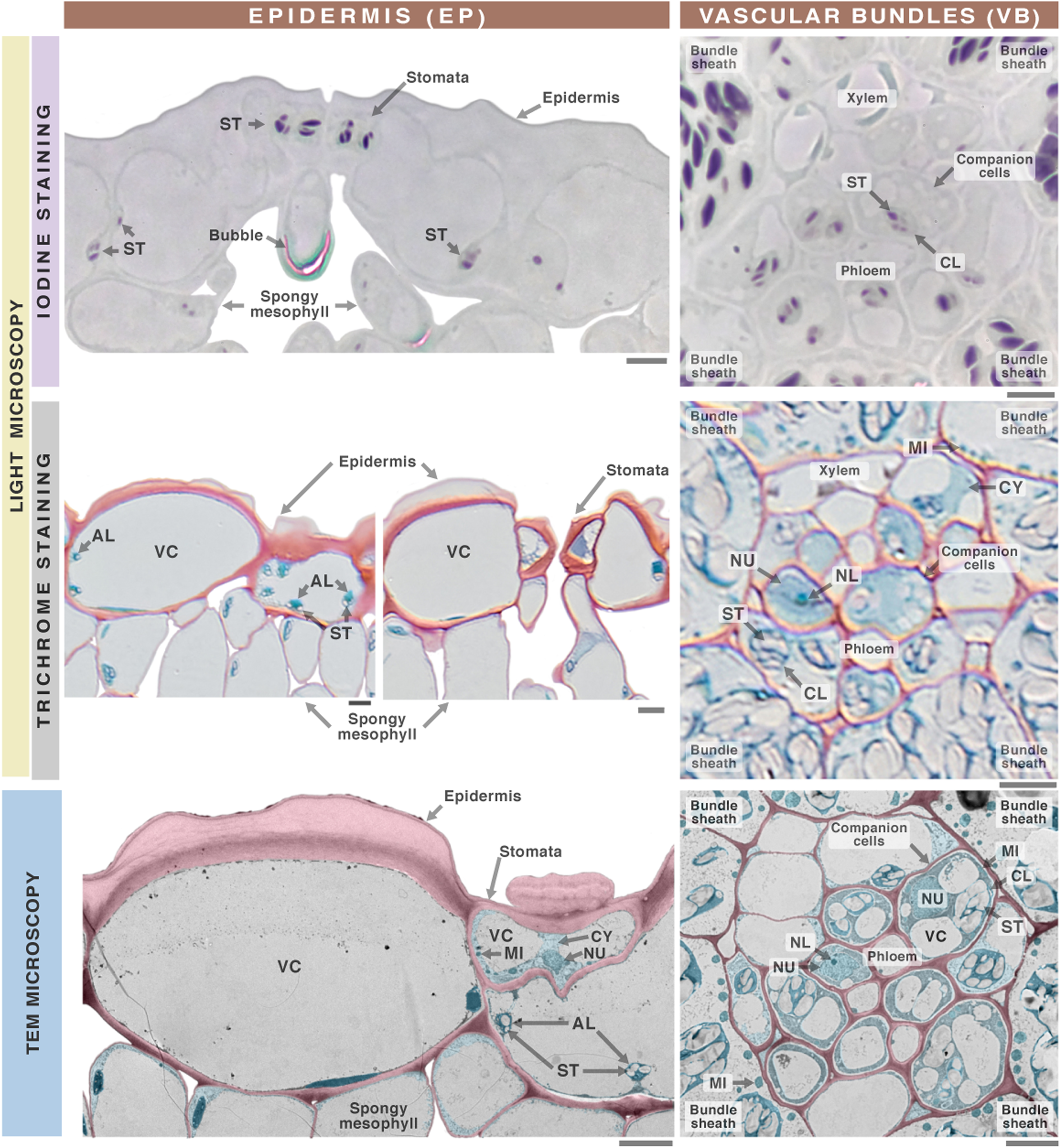
Resin-based microtechnique resolution: epidermis and vascular bundle cells. Epidermal and vascular bundle cells from young (4-week-old) *A. cruentus* leaves were digitally isolated from light and electron micrographs (cells are not the same). Light micrographs display trichrome staining, whereas TEM images are false-colored to match trichrome hues. Vacuole (VC), chloroplast (CL), amyloplast (AL), starch granule (ST), nucleus (NU), nucleolus (NL), cytosol (CY), and mitochondrion (MI) are indicated. High-resolution resin sections revealed numerous ultrastructures confirmed by TEM. The epidermis and vascular bundles comprise distinct cell types: epidermal cells, stomata, and trichomes in the epidermis; xylem, phloem, and diverse accessory cells in the vascular bundles. Epidermal cells contain minimal proteoplasm and typically retain only traces of amyloplasts with starch granules. In contrast, stomatal cells contain organelle-rich proteoplasm similar to mesophyll cells and accumulate abundant starch. Xylem and phloem conducting cells are generally hollow, while their associated accessory cells display proteoplasm enriched with organelles, cytosol, and starch. Vacuoles and chloroplasts—or amyloplasts—are present in nearly all leaf cell types.

**SUPPLEMENTARY FIGURE 10.**
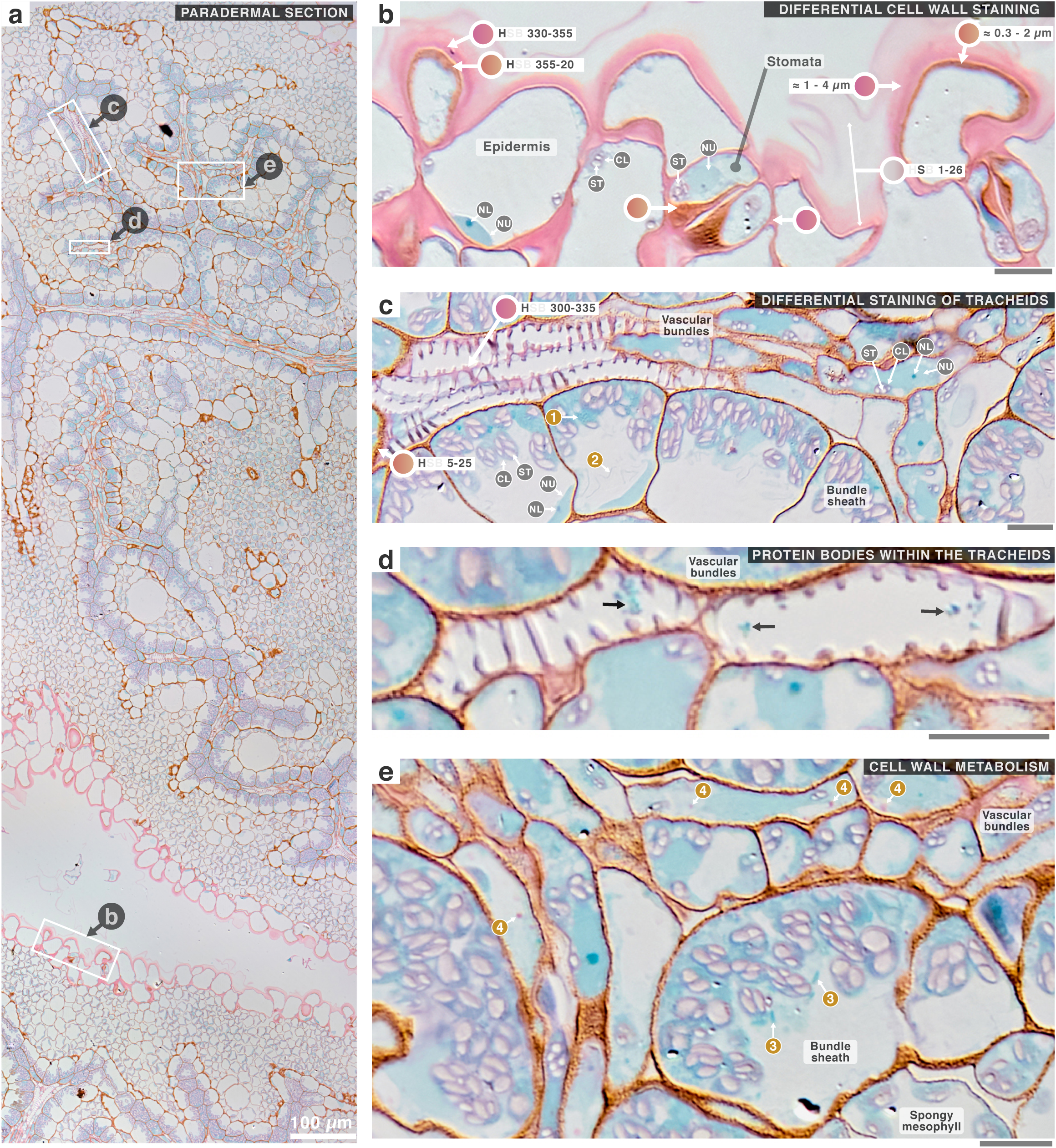
Intense iodine counterstaining reveals cell wall composition and ultrastructures. **a**, Paradermal section of *A. cruentus* leaves stained with trichrome, in which iodine provided an intense counterstain. **b-e**, Magnified regions from the boxed areas are shown (scale bars = 10 µm). Vacuole (VC), chloroplast (CL), starch granule (ST), nucleus (NU), nucleolus (NL), and cytosol (CY) are annotated. The iodine counterstain strongly stained starch granules, produced widespread cell wall hue shifting, and enhanced visualization of ultrastructural detail. Epidermal cell walls exhibited a bilayered organization: an inner orange layer and an outer magenta layer. The outer layer displayed a saturation gradient, transitioning from strongly saturated internally to nearly colorless externally, ending in a sharply saturated boundary. Most cell walls shifted toward orange, while xylem tracheids retained a magenta hue similar to the epidermal outer layer. Occasional protein bodies were visible within the tracheids. Additional ultrastructural features resolved by the iodine contrast included chloroplasts with granular internal texture (**1**), membranes within vacuoles (**2**), rod-shaped organelles (**3**), and vesicular trafficking of wall material (**4**).

**SUPPLEMENTARY FIGURE 11.**
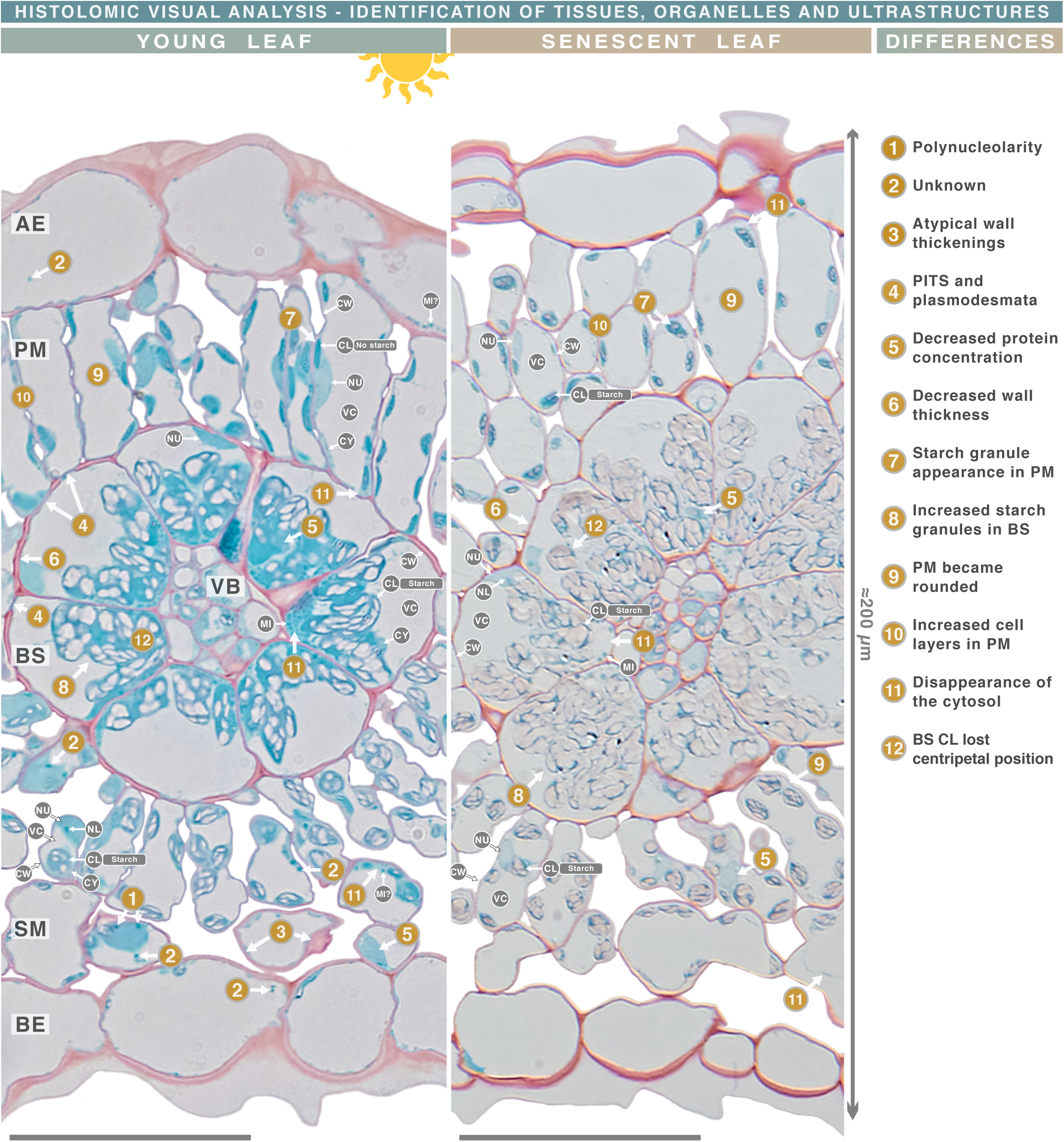
Histolomic analysis – Visual analysis. Light micrographs of cross sections from young (4-week) and senescent (8-week) *A. cruentus* leaves stained with trichrome are shown (scale bar = 50 µm). Adaxial epidermis (AE), palisade mesophyll (PM), bundle sheath (BS), vascular bundles (VB), spongy mesophyll (SM), abaxial epidermis (BE), cell wall (CW), vacuole (VC), chloroplast (CL), starch granule (ST), nucleus (NU), nucleolus (NL), cytosol (CY), and mitochondrion (MI). Visual inspection provides the foundation of histolomic analysis, enabling identification of tissues, general morphology, organelles, and selected ultrastructural features. This step guides segmentation boundaries, feature selection for quantitative analysis, and recognition of unique traits. The amaranth leaf averages ≈200 µm in thickness, comprising 8–10 cells (excluding VB), and shows clear Kranz anatomy and strong differentiation between PM and SM. BS, VB, PM, and SM are organelle-rich, whereas the epidermis contains minimal proteoplasm. Several ultrastructural traits were observed: nuclei with apparent polynucleolarity, unidentified proteinaceous bodies (possibly mitochondria) with atypical size or position, unusual wall thickenings in SM, and PITS-type plasmodesmata between PM and BS. Compared to young leaves, senescent leaves exhibited reduced protein concentration, thinner cell walls, increased starch in PM and BS, rounded PM cells with more layers, disappearance of cytosol, and loss of the centripetal arrangement of BS chloroplasts.

**SUPPLEMENTARY FIGURE 12.**
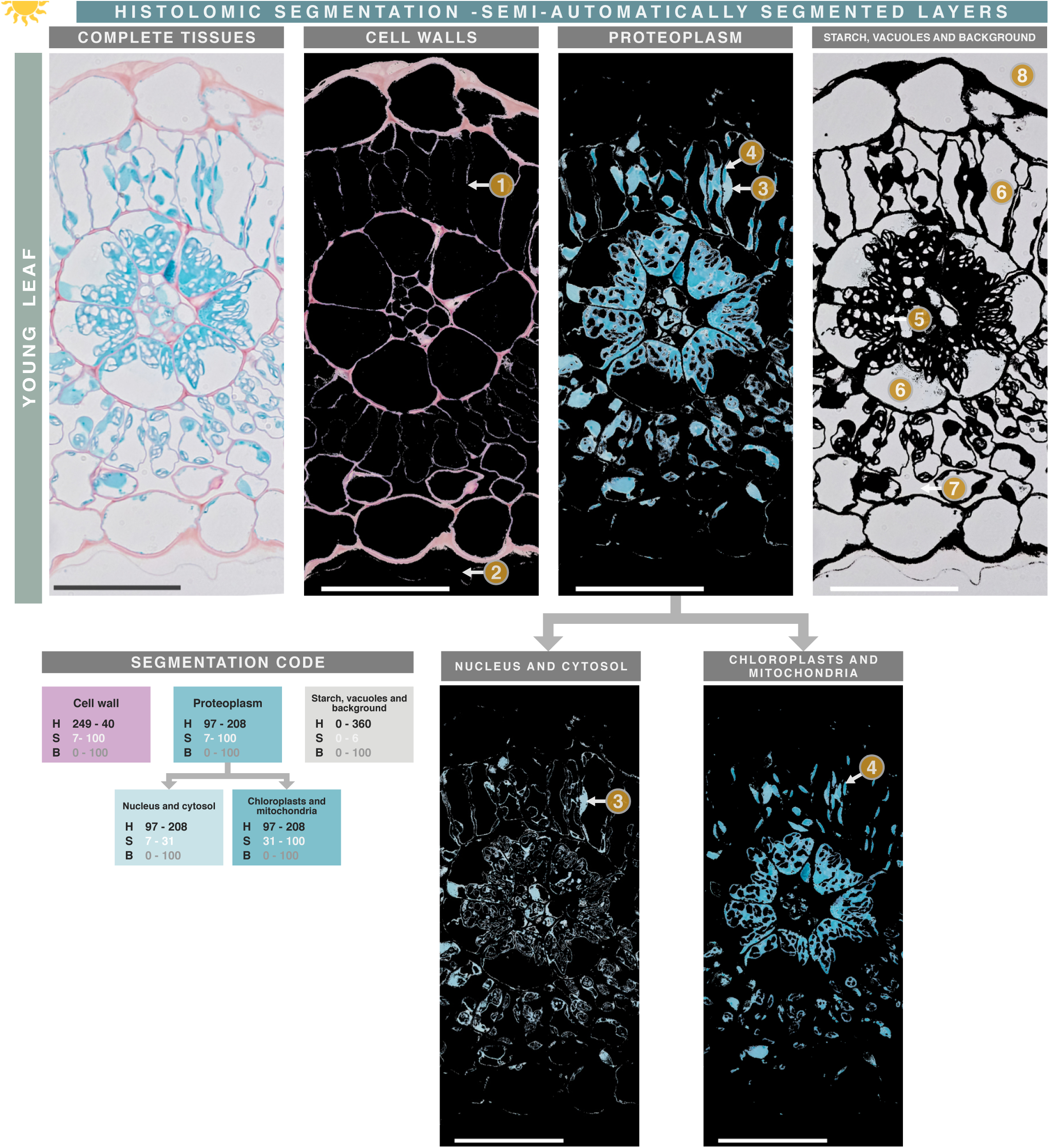
Histolomic analysis – Semi-automated segmentation. A trichrome-stained cross section of a young *A. cruentus* leaf was digitally segmented to isolate tissue components (scale bar = 50 µm). Segmentation was performed in MATLAB using the “Color Threshold” app and custom “segmentation code” settings. Thin or lightly stained mesophyll walls showed discontinuities (**1**), as did the outer epidermal walls (**2**). Variations in proteoplasm saturation allowed visual discrimination of organelles and cytosol, enabling secondary segmentation into low-saturation proteoplasm (**3**, e.g., cytosol, nucleus) and high-saturation proteoplasm (**4**, e.g., chloroplasts, mitochondria). Iodine staining was absent in starch granules in this section (**5**), causing them to be grouped with other unstained regions such as vacuoles (**6**), intercellular air spaces (**7**), and resin background (**8**). Semi-automated segmentation is efficient but struggles with thin or weakly stained walls and requires subsequent manual refinement to subdivide layers by tissue type.

**SUPPLEMENTARY FIGURE 13.**
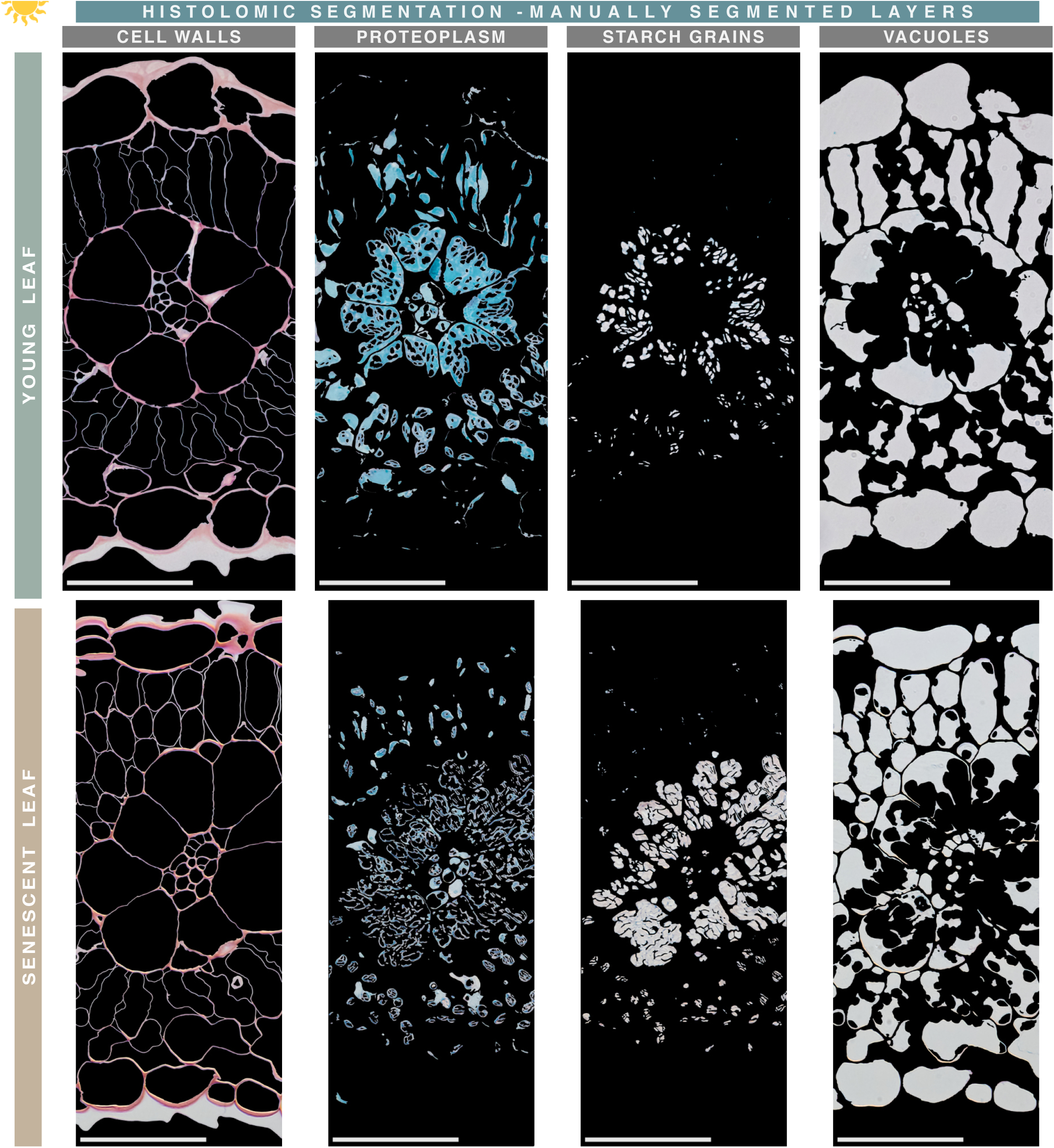
Histolomic analysis – Manual segmentation of tissue components of an entire section. Manually segmented layers from light micrographs of trichrome-stained cross sections of young and senescent *A. cruentus* leaves (scale bar = 50 µm). Manual segmentation is time-consuming but essential for delineating thin or weakly stained cell walls that are lost during semi-automated segmentation. Each segmented layer represents more than a visually appealing image: it contains quantitative information—size, perimeter, area, shape—for morphometric analysis, as well as color-intensity data (densitometry) for compositional measurements. Clear, continuous cell boundaries are critical for accurate morphometric analysis. Manual segmentation is also necessary to separate layers by tissue type for independent study.

**SUPPLEMENTARY FIGURE 14.**
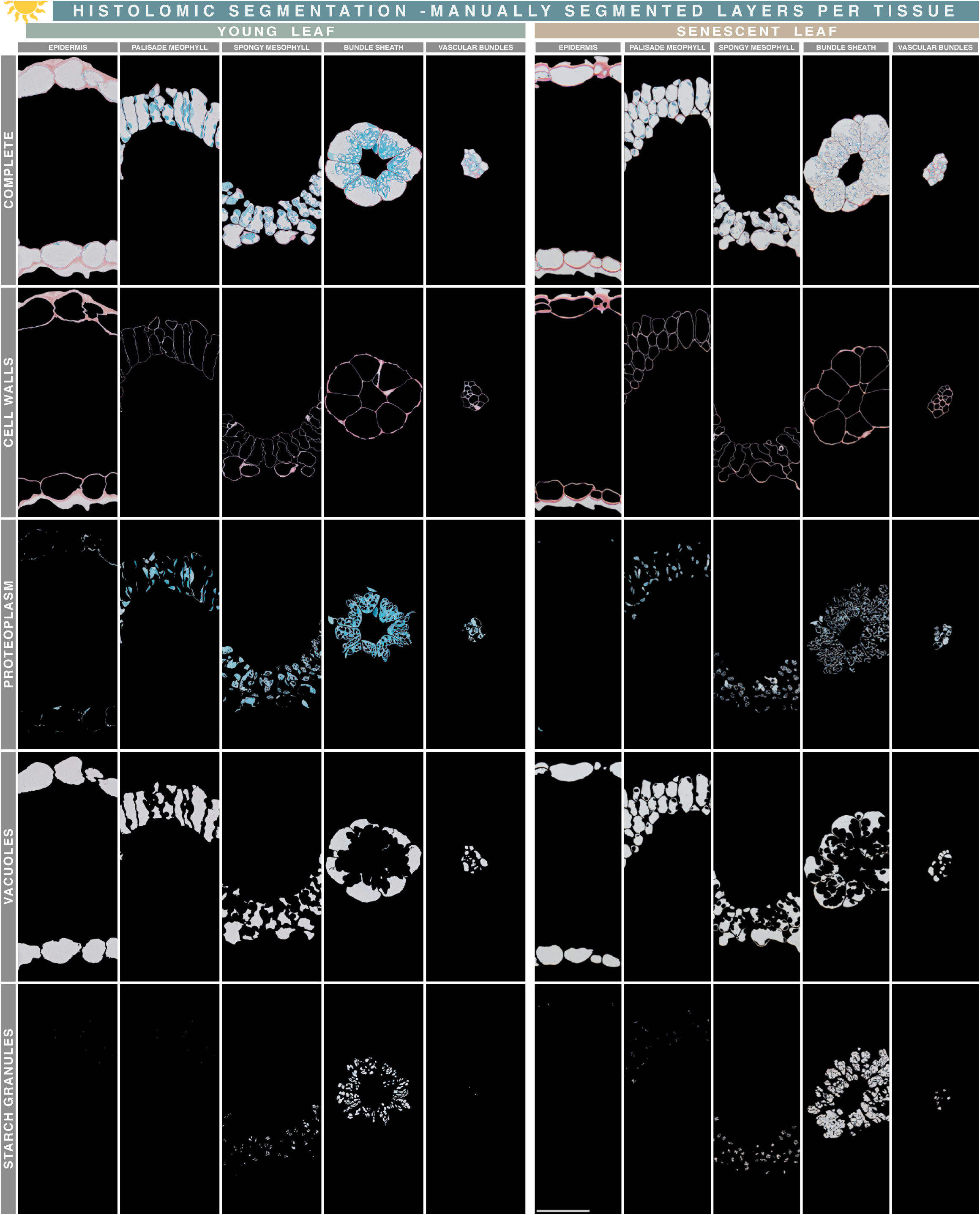
Histolomic analysis – Manual segmentation of t issue components per t issue. Manually segmented layers from light micrographs of trichrome-stained cross sections of young and senescent *A. cruentus* leaves (scale bar = 50 µm). The aim of histolomics is the individualized phenotypic characterization of each tissue within an organ. We envision cell-by-cell segmentation and analysis as the long-term goal of histolomics, although such work is presently very labor-intensive when performed manually.

**SUPPLEMENTARY FIGURE 15.**
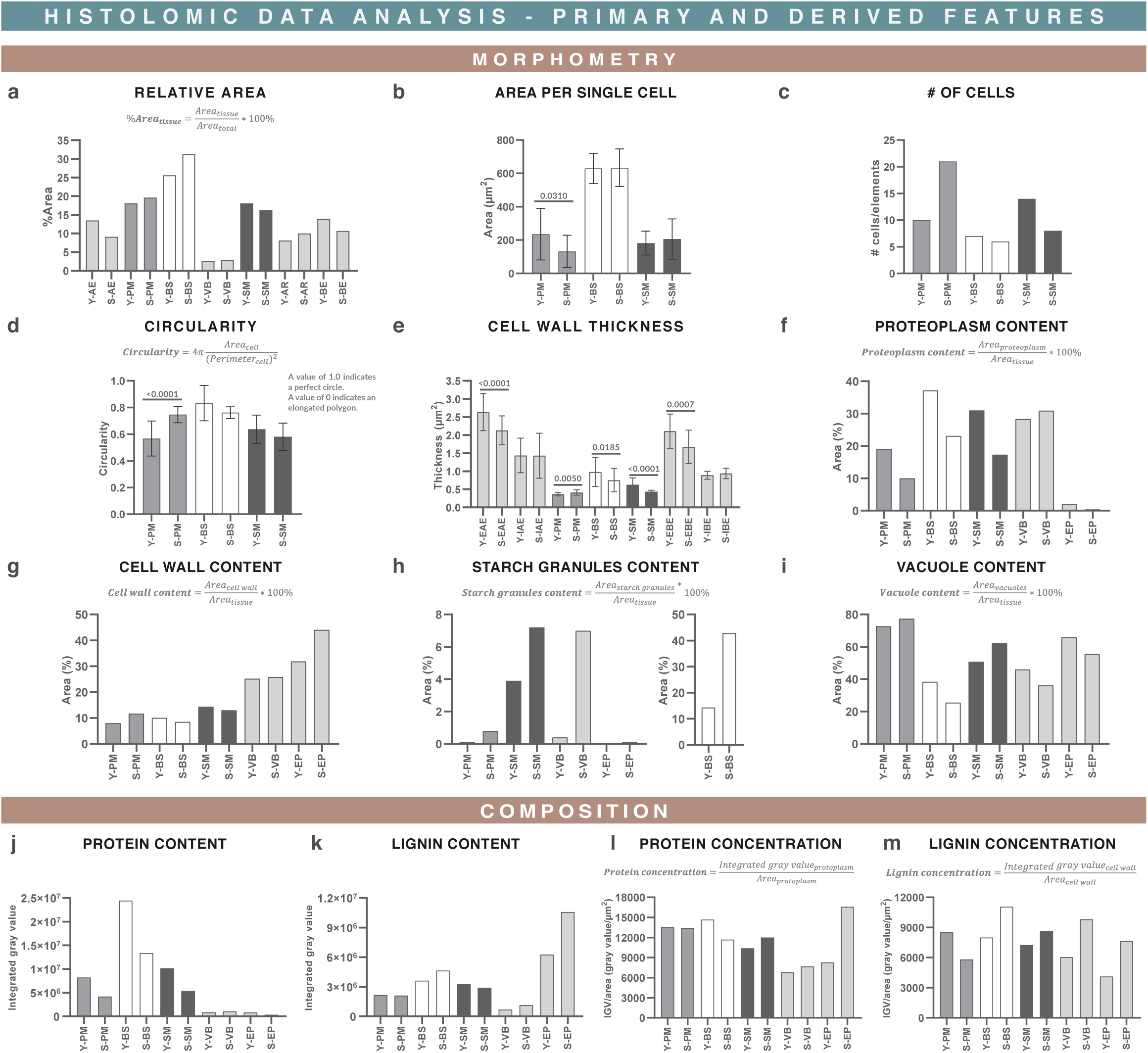
Histolomic analysis – Data analysis. Morphometric and compositional analysis of trichrome-stained cross sections of young and senescent *A. cruentus* leaves. Tissue abbreviations: adaxial epidermis (AE), abaxial epidermis (BE), adaxial inner epidermis (IAE), adaxial outer epidermis (EAE), abaxial inner epidermis (IBE), abaxial outer epidermis (EBE), palisade mesophyll (PM), spongy mesophyll (SM), bundle sheath (BS), vascular bundles (VB), air spaces (AR), young leaf (Y), and senescent leaf (S). For replicated measurements, t-tests comparing young vs. senescent leaves were performed; differences are shown at p < 0.05. Histolomics requires its own set of features and formulas to characterize morphometric and compositional traits beyond visual inspection. We propose three categories of histolomic features: primary, derived, and composite. Primary features include direct measurements such as cell count, thickness, area, perimeter, and total protein or lignin content. Derived features are ratios or combinations of primary features (e.g., protein or lignin concentrations, organelle content). Composite features combine multiple primary and derived features, can include weighting, and allow quantitative comparisons of complex tissue behavior (Supplementary Fig. 24). This figure presents primary and derived features and their formulas (when applicable). Their values revealed quantitative differences in senescent compared with young leaves, consistent with visual observations: reduced protein concentration (**l**), decreased cell wall thickness (**e**), increased starch granules in PM (**h**), rounded PM cells (**d**), and additional PM layers (**c**). Kranz-related traits were also revealed: BS was by far the largest tissue (**a**), had the largest cells (**b**), contained less vacuole space (**i**), had the greatest proteoplasm content (**f**), and accumulated the most protein **(j**) and starch granules (**h**). BS also showed slightly higher lignin concentration (**m**) and thicker cell walls (**g**, **e**). Protein and lignin measurements are semi-quantitative because they were not calibrated to absolute standards (**j**–**m**); however, in comparative analyses, relative differences are typically more informative than absolute values.

**SUPPLEMENTARY FIGURE 16.**
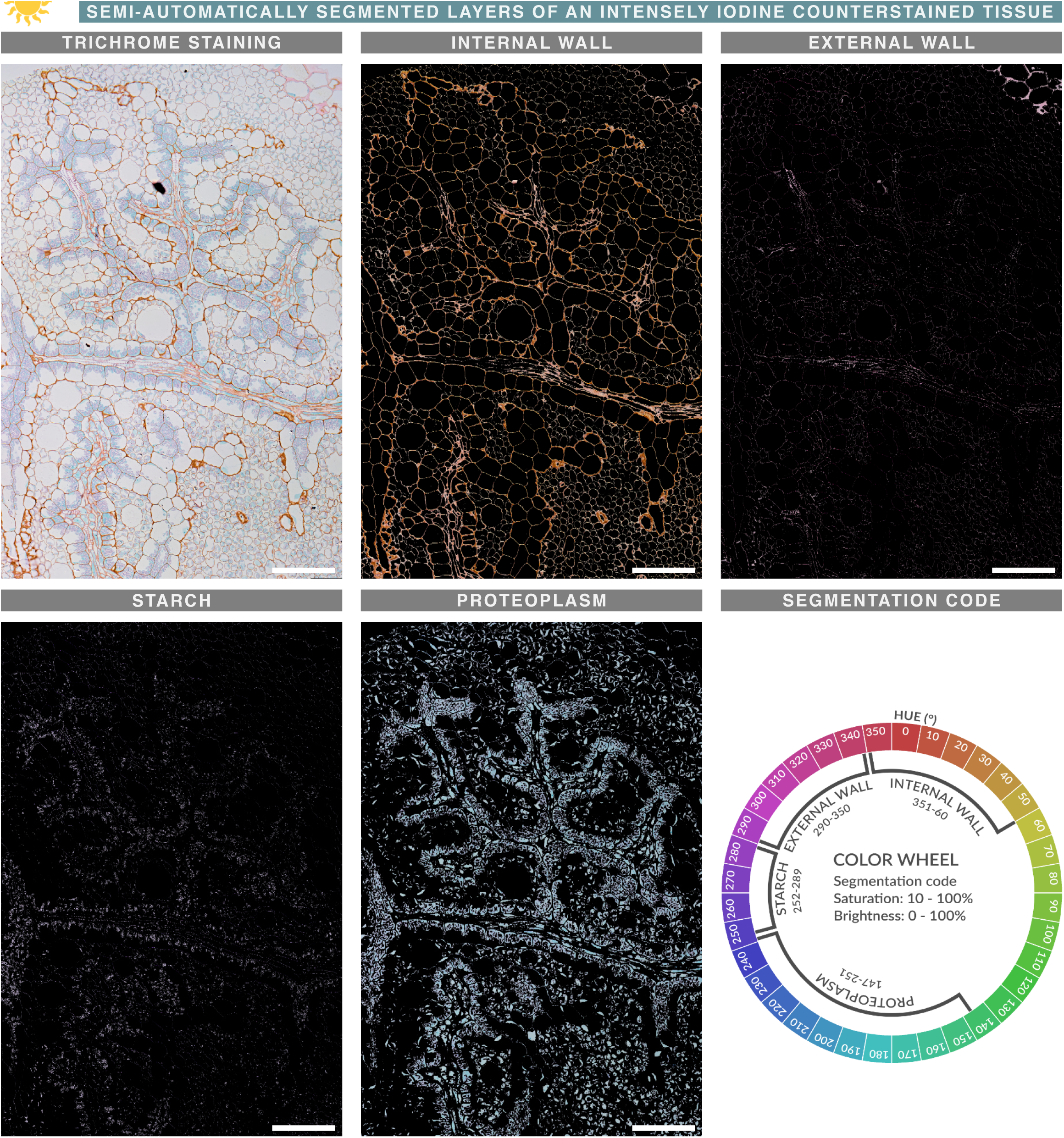
Intense iodine counterstaining allows further color (composition) segmentation. Semi-automatic color segmentation of a trichrome-stained paradermal section of an *A. cruentus* leaf (scale bar = 100 µm). The color wheel illustrates the hue ranges assigned to each component and the saturation/brightness thresholds used for segmentation. Intense iodine counterstaining shifted nearly all cell walls toward orange, except for the epidermal walls, which exhibited an inner orange layer and an outer magenta layer. The inner layer displayed a broad range of reddish, orange, and yellow hues, whereas the outer layer showed magenta to reddish hues. Color segmentation revealed that although most walls shifted toward orange, they still retained magenta tones, suggesting the presence of two compositionally distinct wall layers. Intense iodine counterstaining also enabled segmentation of starch granules and did not produce major hue shifts in the proteoplasm, unlike its effects on cell walls. However, the starch granule hue range overlapped slightly with that of the epidermal outer wall, making these layers “impure.” Despite this, intense iodine counterstaining enhanced the contrast of thin mesophyll walls and expanded the detectable information on cell wall composition, thereby improving boundary identification.

**SUPPLEMENTARY FIGURE 17.**
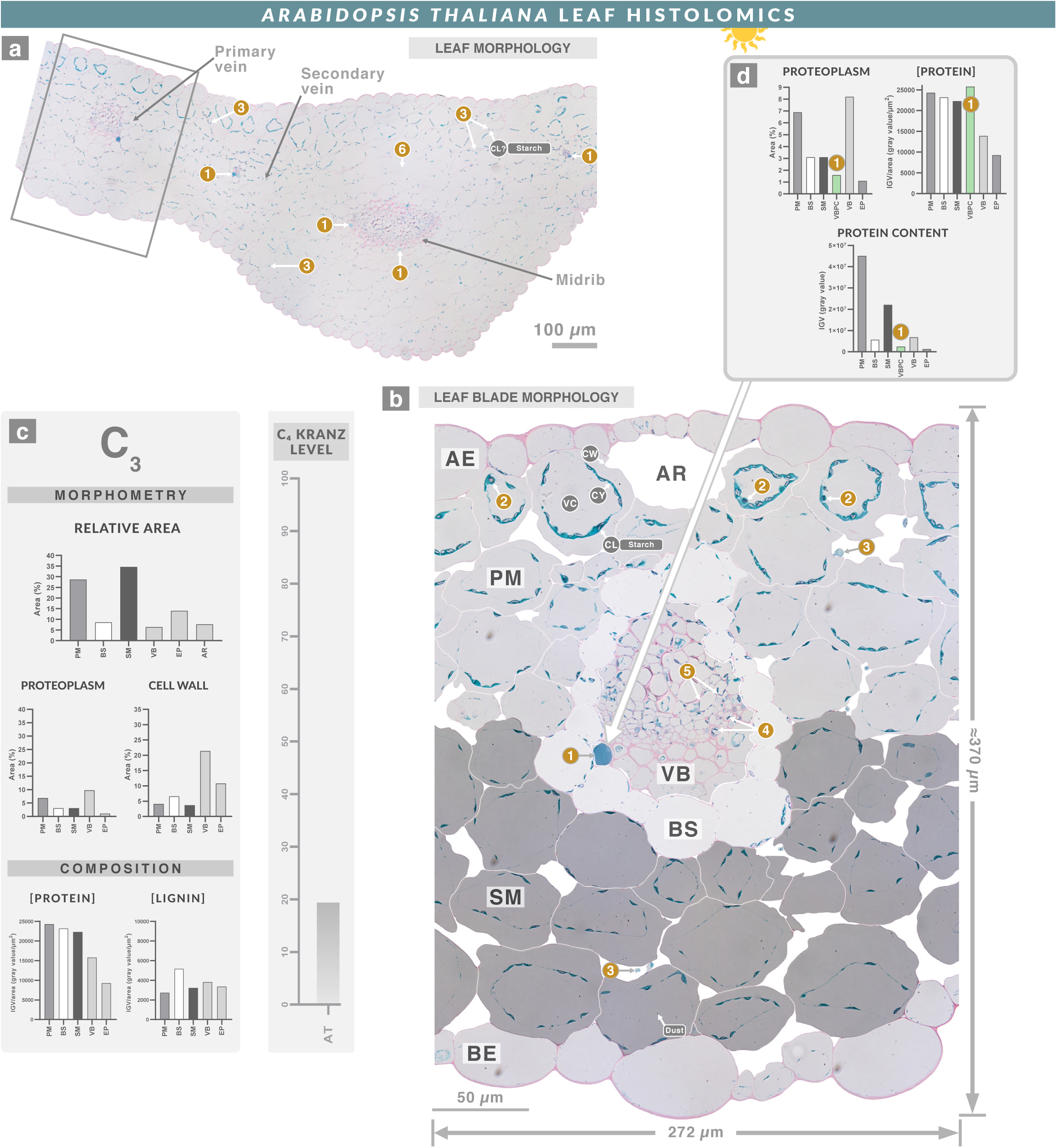
Histolome of *Arabidopsis thaliana* leaf. **a**, Trichrome-stained cross section of the midrib of a *Arabidopsis thaliana* leaf. **b**, Magnification of the boxed area in **a**. **c,** Derived (morphometry and composition) and composite (C₄ kranz level) features of the tissue (same scales for all plant species). **d**, Primary features of the vascular bundle proteinaceous cell (VBPC). Adaxial epidermis (AE), abaxial epidermis (BE), palisade mesophyll (PM), spongy mesophyll (SM), bundle sheath (BS), vascular bundle (VB), air spaces (AR). Cell wall (CW), vacuole (VC), chloroplast (CL), starch granule (ST), nucleus (NU), nucleolus (NL), cytosol (CY), and mitochondrion (MI). Although not all plants have a developed BS or differentiated PM and SM, these tissues are identified here for comparative purposes. Brightness was increased in BS and decreased in SM to visually distinguish them.The A. thaliana leaf had a small BS tightly surrounded by two PM layers and three SM layers, lacking a wreath-shaped arrangement. All cells showed low proteoplasmic content (**c**), except for a distinct VBPC in the VB, characterized by smooth, well-nourished proteoplasm occupying nearly the entire cytoplasm (**1**). One VBPC was present in the primary vein, and two were observed in the secondary vein and midrib, though with lower proteoplasm and protein concentration than in the primary vein (**a**). VBPCs contained up to 26% of the VB protein content in the primary vein (**d**). Additional ultrastructural observations included organelles with purple granules (**2**), extracellular proteoplasm (**3**), dark purple rod-shaped organelles in the VB (**4**), intermittent staining in VB cell walls (**5**), and apparently ruptured cells (**6**).

**SUPPLEMENTARY FIGURE 18.**
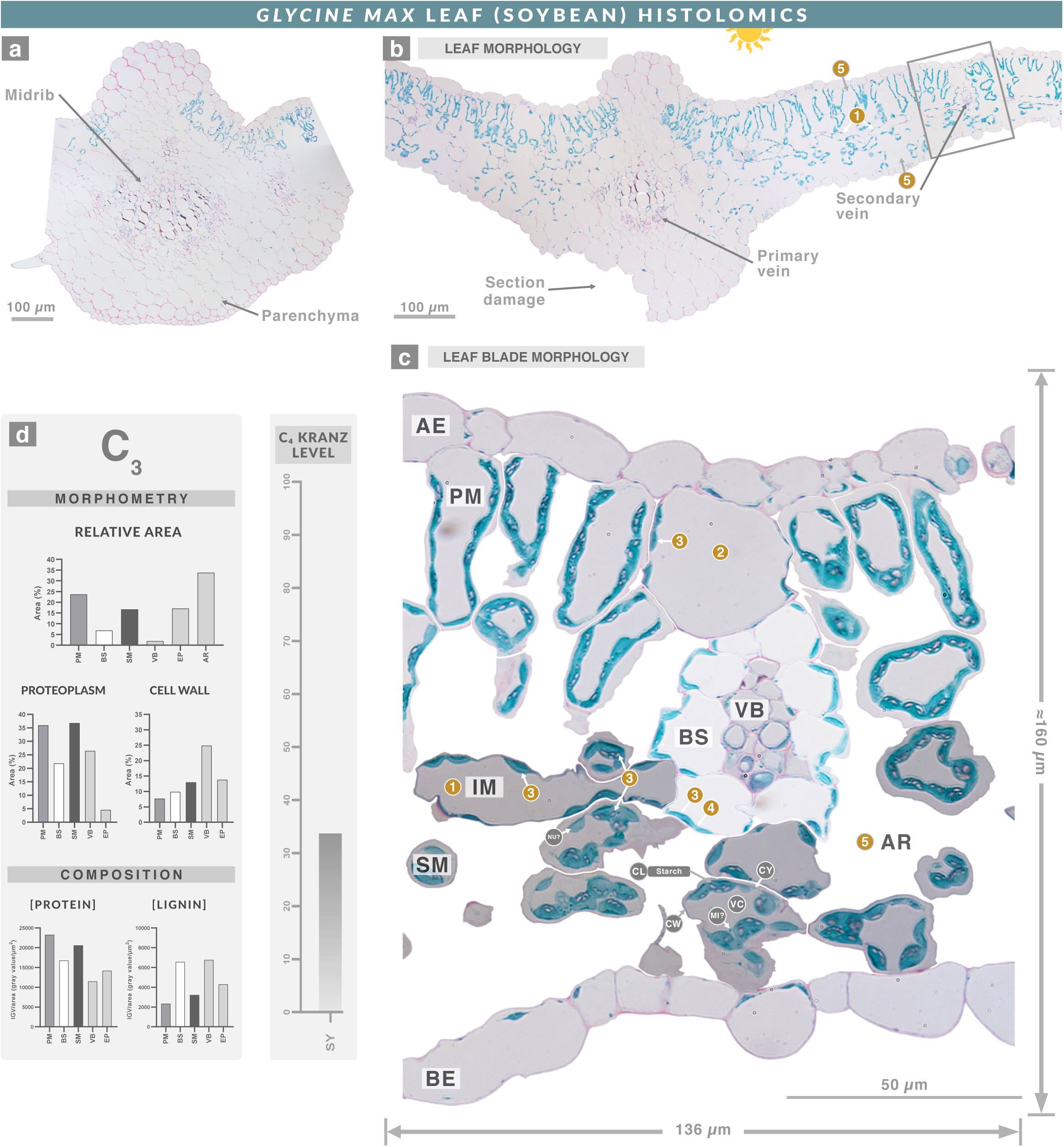
Histolome of *Glycine max* leaf. **a**, Trichrome-stained cross section of the midrib of a *Glycine max* leaf leaf. **b**, Cross section of the leaf blade. **c**, Magnification of the boxed area in **b**. **d**, Derived (morphometry and composition) and composite (C₄ kranz level) features of the tissue (same scales for all plant species). The *G. max* leaf had a small BS loosely surrounded by one PM layer and two SM layers, thus not forming a wreath-shaped structure. A distinct rod-shaped mesophyll cell type, arranged perpendicular to the PM, was observed and termed intermediate mesophyll (IM, **1**). IM was present throughout the blade, along with other rounded PM cells (**2**). Compositionally, IM did not accumulate starch granules like BS and (**2**), in contrast to PM and SM (**3**). Although *G. max* is a C₃ species, BS chloroplasts displayed centrifugal orientation—a feature typically associated with C₄ plants (**4**). The leaf also contained abundant intercellular air spaces (**5**).

**SUPPLEMENTARY FIGURE 19.**
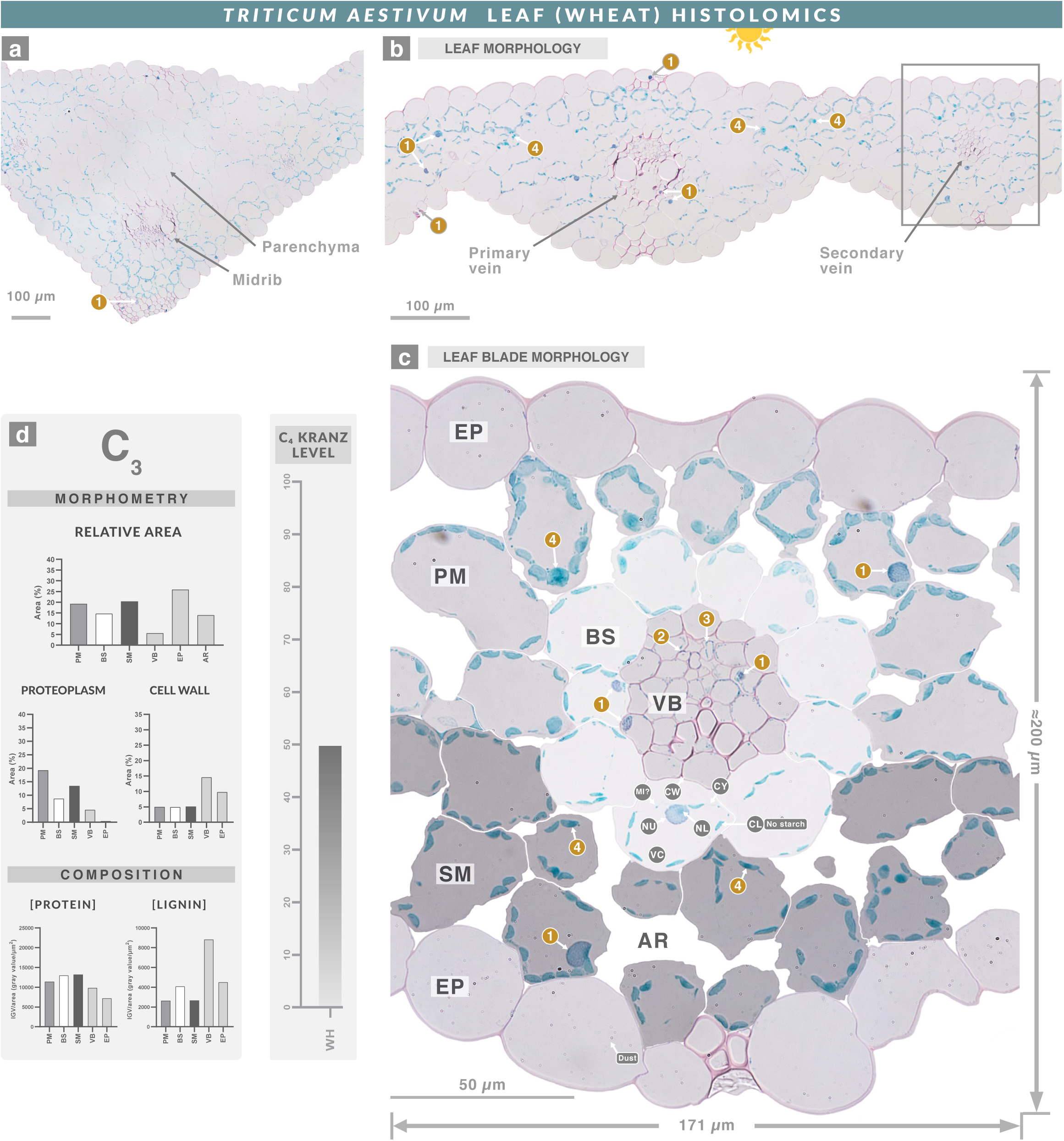
Histolome of Triticum aestivum leaf. **a**, Trichrome-stained cross section of the midrib of a *Triticum aestivum* leaf. **b**, Cross section of the leaf blade. **c**, Magnification of the boxed area in **b**. **d**, Derived (morphometry and composition) and composite (C₄ kranz level) features of the tissue (same scales for all plant species). The *T. aestivum* leaf had a large BS tightly surrounded by one layer of undifferentiated PM and two layers of SM, forming a wreath-shaped arrangement. Additional notable features included magenta granules within turquoise-stained nuclei (**1**), intermittent magenta-purple staining along VB cell walls (**2**), minimal proteoplasm in the VB (**3**), and polarized protein distribution within certain organelles (**4**).

**SUPPLEMENTARY FIGURE 20.**
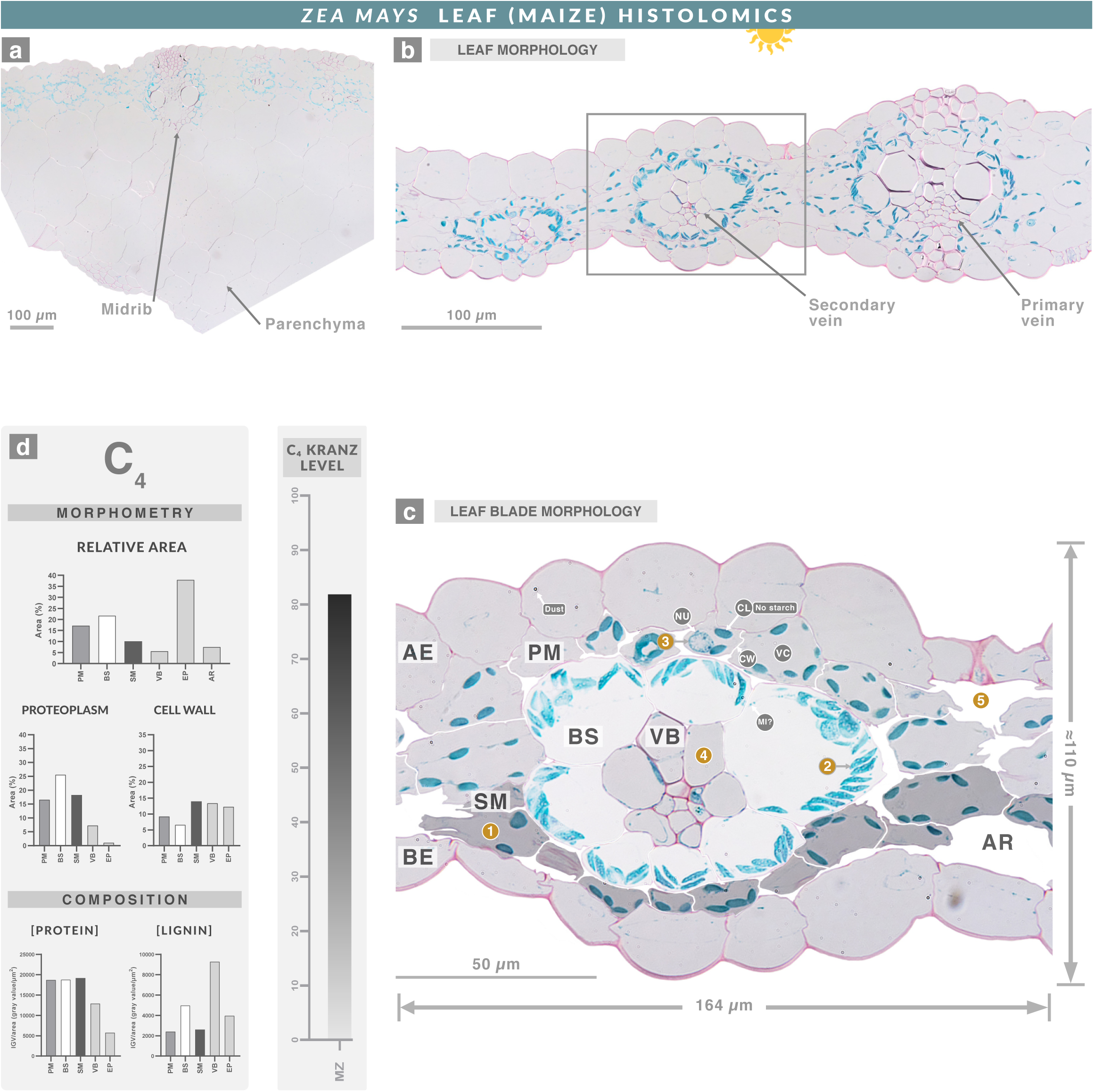
Histolome of *Zea mays* leaf. **a**, Trichrome-stained cross section of the midrib of a *Z. mays* leaf. **b**, Cross section of the leaf blade. **c**, Magnification of the boxed area in **b**. **d**, Derived (morphometry and composition) and composite (C₄ kranz level) features of the tissue (same scales for all plant species). The *Z. mays* leaf featured a very large BS tightly surrounded by one undifferentiated PM layer and one SM layer, resulting in a wreath-shaped anatomy. SM cells between VB were elongated with sharp edges (**1**), and BS chloroplasts were centripetally positioned (**2**). Additional observations included granulated nuclei (**3**), low proteoplasmic content in the VB (**4**), and very few intercellular air spaces (**5**).

**SUPPLEMENTARY FIGURE 21.**
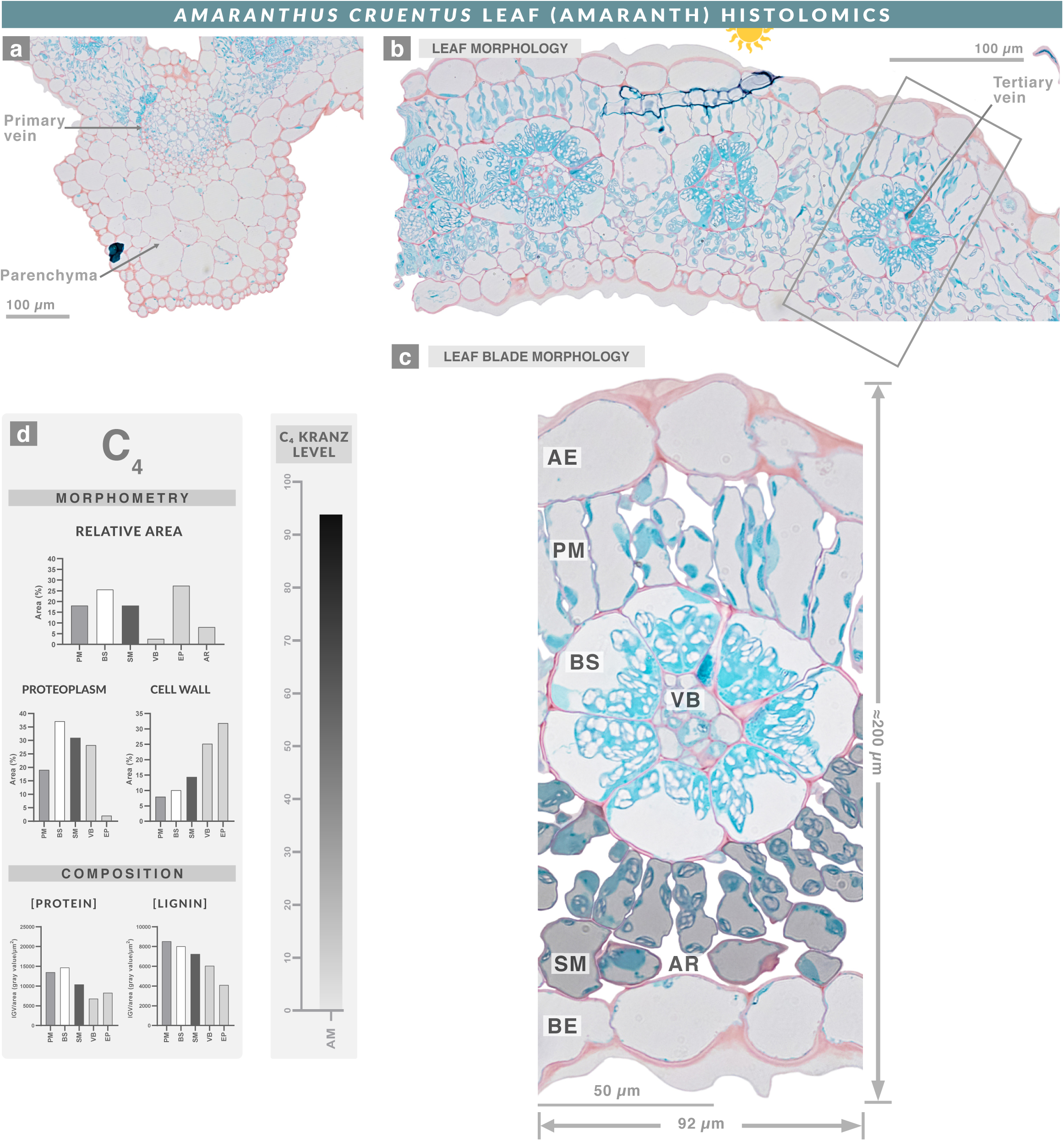
Histolome of *Amaranthus cruentus* leaf. **a**, Trichrome-stained cross section of a *A. cruentus* leaf midrib. **b**, Cross section of the leaf blade. **c**, Magnification of the boxed area in **b**. **d**, Derived (morphometry and composition) and composite (C₄ kranz level) features of the tissue (same scales for all plant species). The *A. cruentus* leaf had a large BS tightly surrounded by one layer of differentiated PM and two layers of SM, forming a wreath-shaped structure. Additional details are provided in Supplementary Figure 11.

**SUPPLEMENTARY FIGURE 22.**
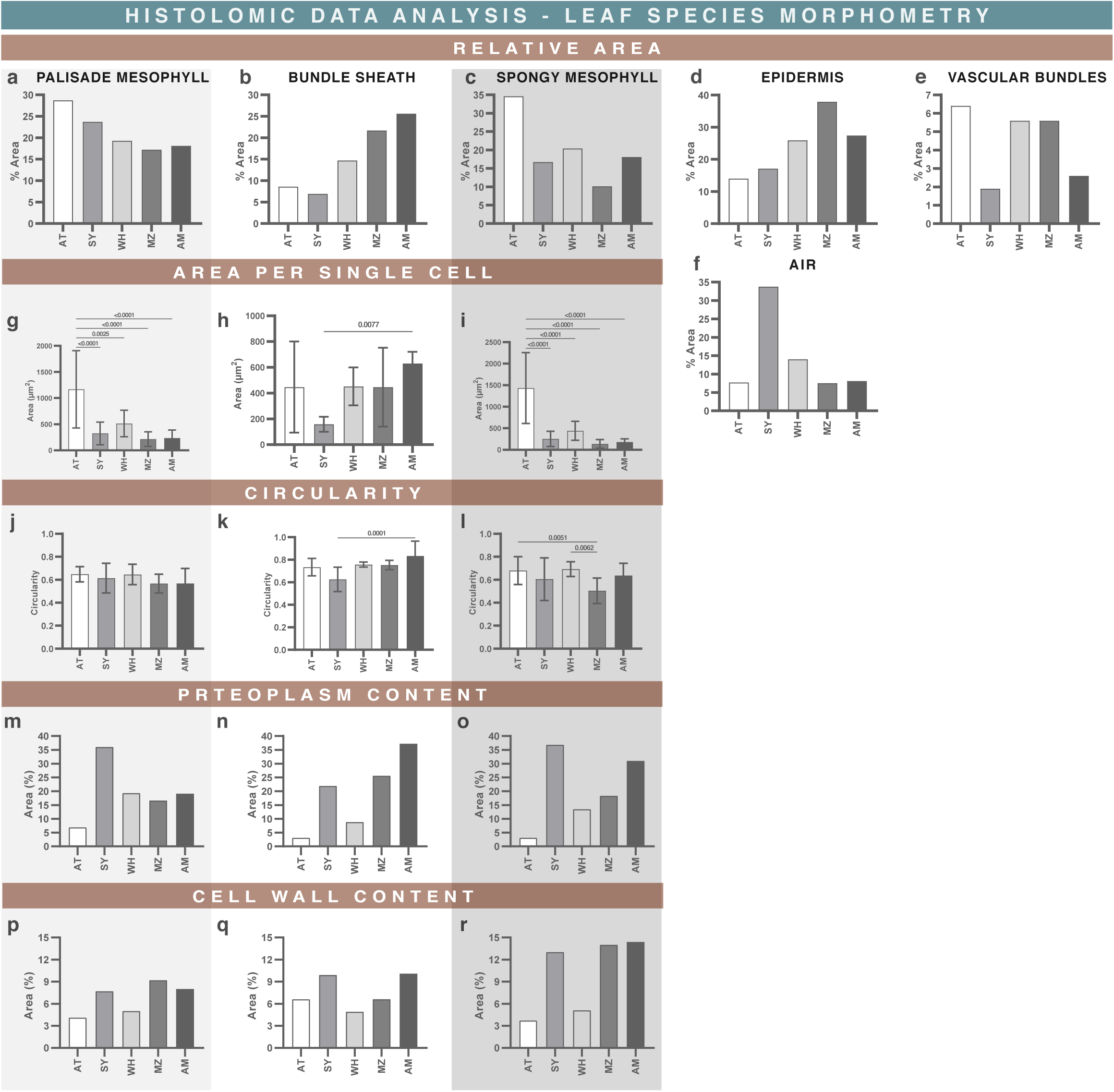
Histolomic morphometry analysis of different plant leaves. Morphometric analyses of *A. thaliana* (AT), *G. max* (SY), *T. aestivum* (WH), *Z. mays* (MZ), and *A. cruentus* (AM) leaves. For measurements with replicates, a one-way ANOVA with Tukey’s post-test was performed, with significant values (p < 0.01) indicated. Species are arranged from left to right according to ascending C₄ kranz level (AT → SY → WH → MZ → AM), with the leftmost representing the “most C₃” and the rightmost the “most C₄.” Special emphasis was placed on the primarily photosynthetic tissues—PM, SM, and BS—because they define the wreath shape and overall C₄ Kranz anatomy. Several trends associated with increasing C₄ anatomical specialization were observed: BS area increased (**b**) and its proteoplasm content was higher (**n**), whereas PM area decreased (**a**) and its proteoplasm content declined (**m**). SM showed no clear morphometric trend; its area tended to decrease (**c**), while its proteoplasm content tended to increase (**o**). Although thicker BS cell walls are characteristic of C₄ mesophyll anatomy, no consistent trend was detected across species (**p**, **q**, **r**). The epidermis displayed an unexpected trend of increasing size, interrupted only by MZ (**d**). Vascular bundles exhibited no clear pattern. Circularity did not reliably capture evident differences in tissue shape—no significant differences were detected between some PM and SM despite obvious disparities in micrographs (**j**, **l**)—although circularity did distinguish BS of AM and SY, which share a similar shell-like morphology **(k**). Air space content showed no consistent trend (**f**), despite expectations of reduction in C₄ species due to more efficient carbon fixation. Finally, AT exhibited the greatest leaf thickness, number of cell layers, and cell sizes (**g**, **h**, **i**), yet had the lowest proteoplasm content (**m**, **n**, **o**).

**SUPPLEMENTARY FIGURE 23.**
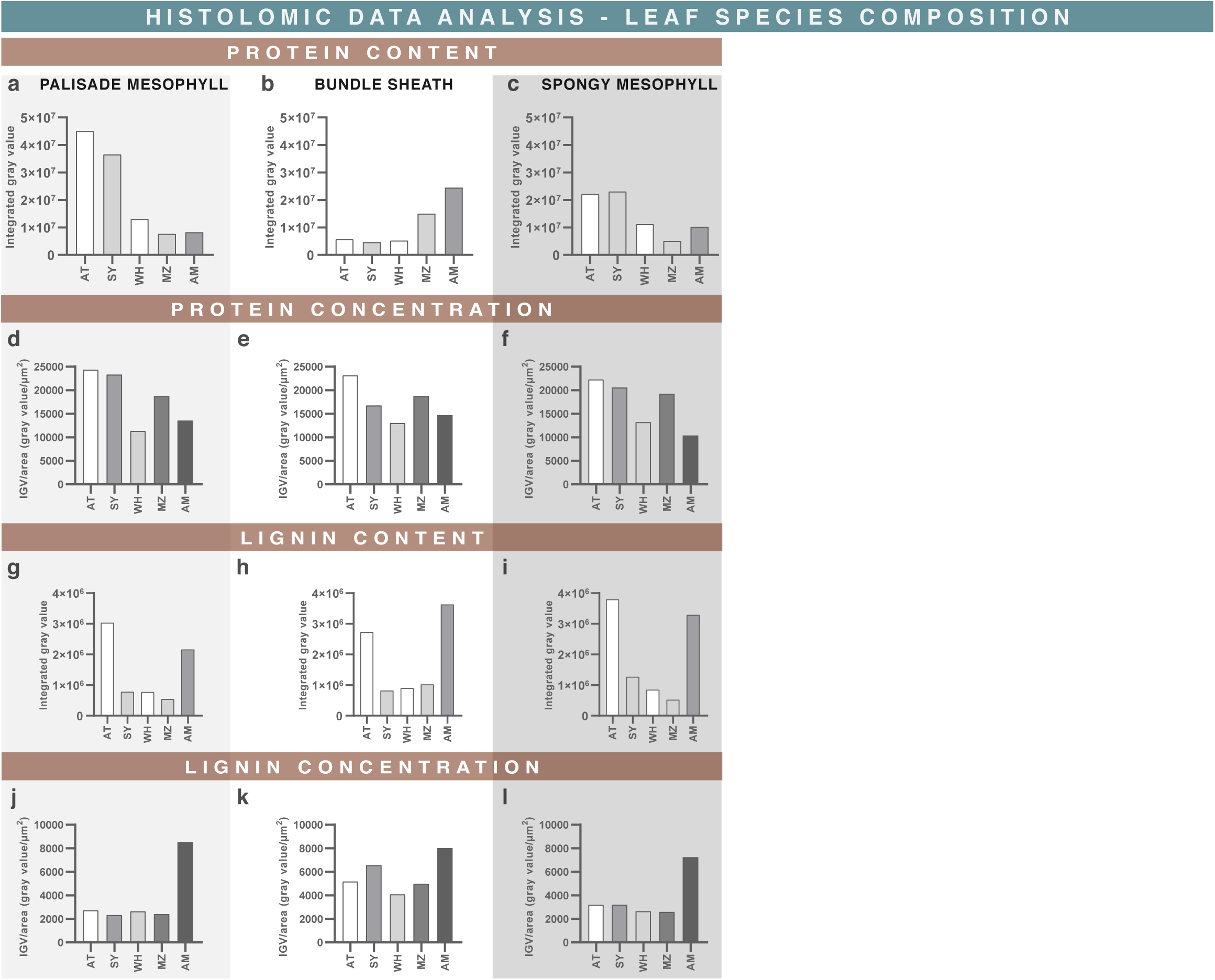
Histolomic compositional analysis of different plant leaves. Compositional analyses of *A. thaliana* (AT), *G. max* (SY), *T. aestivum* (WH), *Z. mays* (MZ), and *A. cruentus* (AM) leaves are presented, again ordered by ascending C₄ kranz level (AT → SY → WH → MZ → AM), with the leftmost representing the “most C₃” and the rightmost the “most C₄.” As in the morphometric analysis, emphasis was placed on the photosynthetic tissues PM, SM, and BS. Trends consistent with increasing C₄ anatomy were observed: protein content decreased in PM and SM (**a**, **c**) and increased in BS (**b**). However, protein concentration showed no consistent pattern across species (**d**–**f**). Lignin content and concentration also exhibited no clear trend (**g**–**l**), although AM tissues consistently had the highest lignin concentration.

**SUPPLEMENTARY FIGURE 24.**
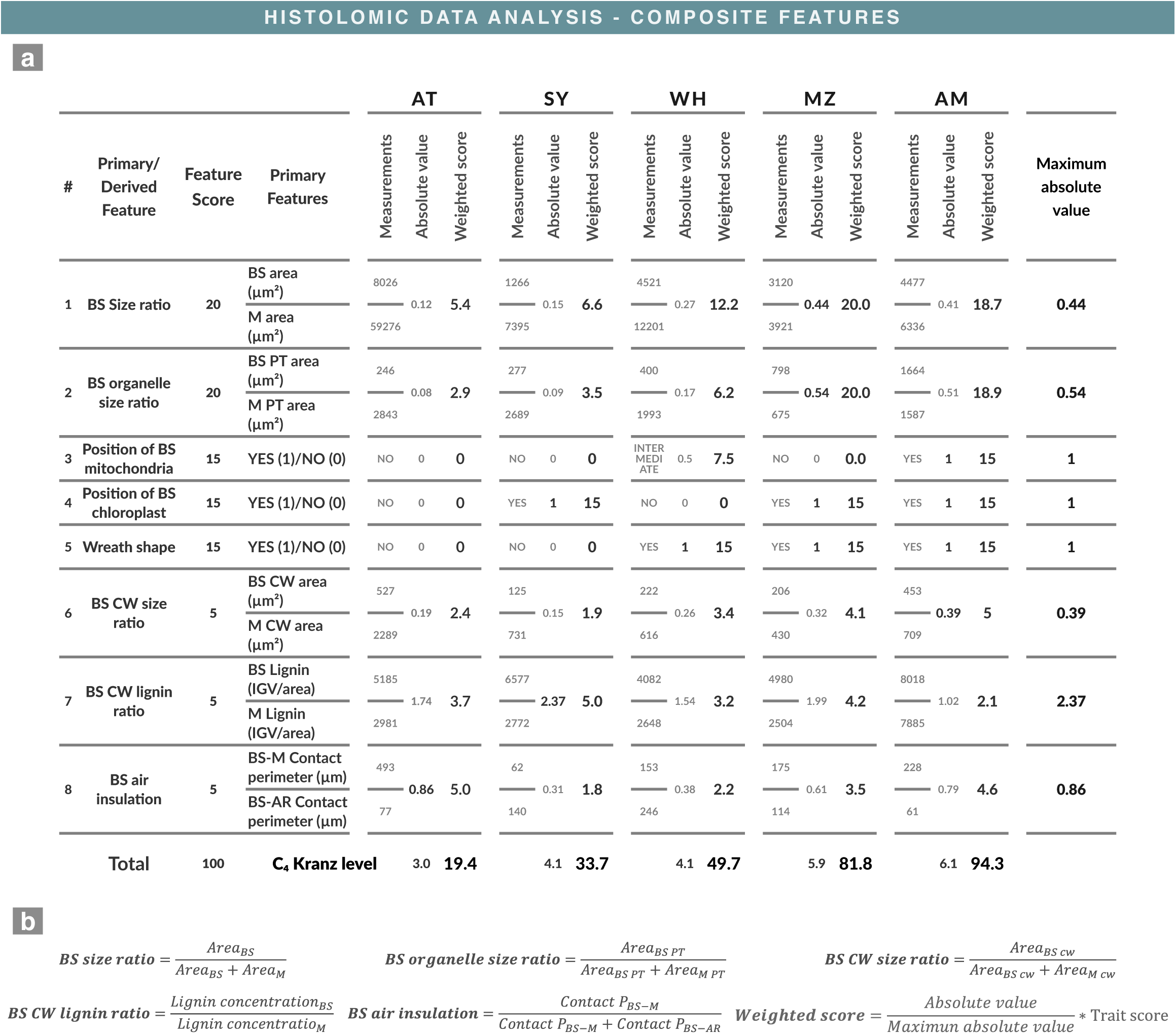
Calculation of the C₄ kranz anatomy level. The C₄ Kranz level of *A. thaliana* (AT), *G. max* (SY), *T. aestivum* (WH), *Z. mays* (MZ), and *A. cruentu*s (AM) leaves was calculated (**a**), and the formulas used are shown (**b**). Bundle sheath (BS), mesophyll (M), cell wall (CW), proteoplasm (PT), air spaces (AR), mitochondrion (MI), chloroplast (CL), and integrated gray value (IGV). Because individual comparisons of primary and derived features did not yield clear conclusions, these features were integrated into a single composite value enabling simple and quantitative comparisons. Primary features describe characteristics of each tissue, such as BS/mitochondria/chloroplast position—indicating whether mitochondria and chloroplasts are centripetally or centrifugally located in BS—and the wreath shape, defined by whether the BS forms a large ring tightly surrounded by one PM layer and one or two SM (or PM) layers. Derived features compare primary traits between tissues, such as the BS size ratio relative to mesophyll area, or analogous comparisons for organelle size, cell wall thickness, lignin concentration, and air insulation. Collectively, this set of morphometric and compositional features comprises the C₄ kranz anatomy. Because each feature contributes with different biological relevance, we assigned weighted scores and calculated a weighted sum that reflects the overall “C₄ kranz level,” a composite histolomic feature. Importantly, this metric is intended as an example of a histolomic application and is not meant to represent a complete description of C₄ photosynthesis. True C₄ identity also involves specific localization of key enzymes—such as glycine decarboxylase complex in BS mitochondria, PEP carboxylase in mesophyll cytosol, and RuBisCO and NADP-malic enzyme in BS chloroplasts—which are not included in the weighting scheme.

**SUPPLEMENTARY FIGURE 25.**
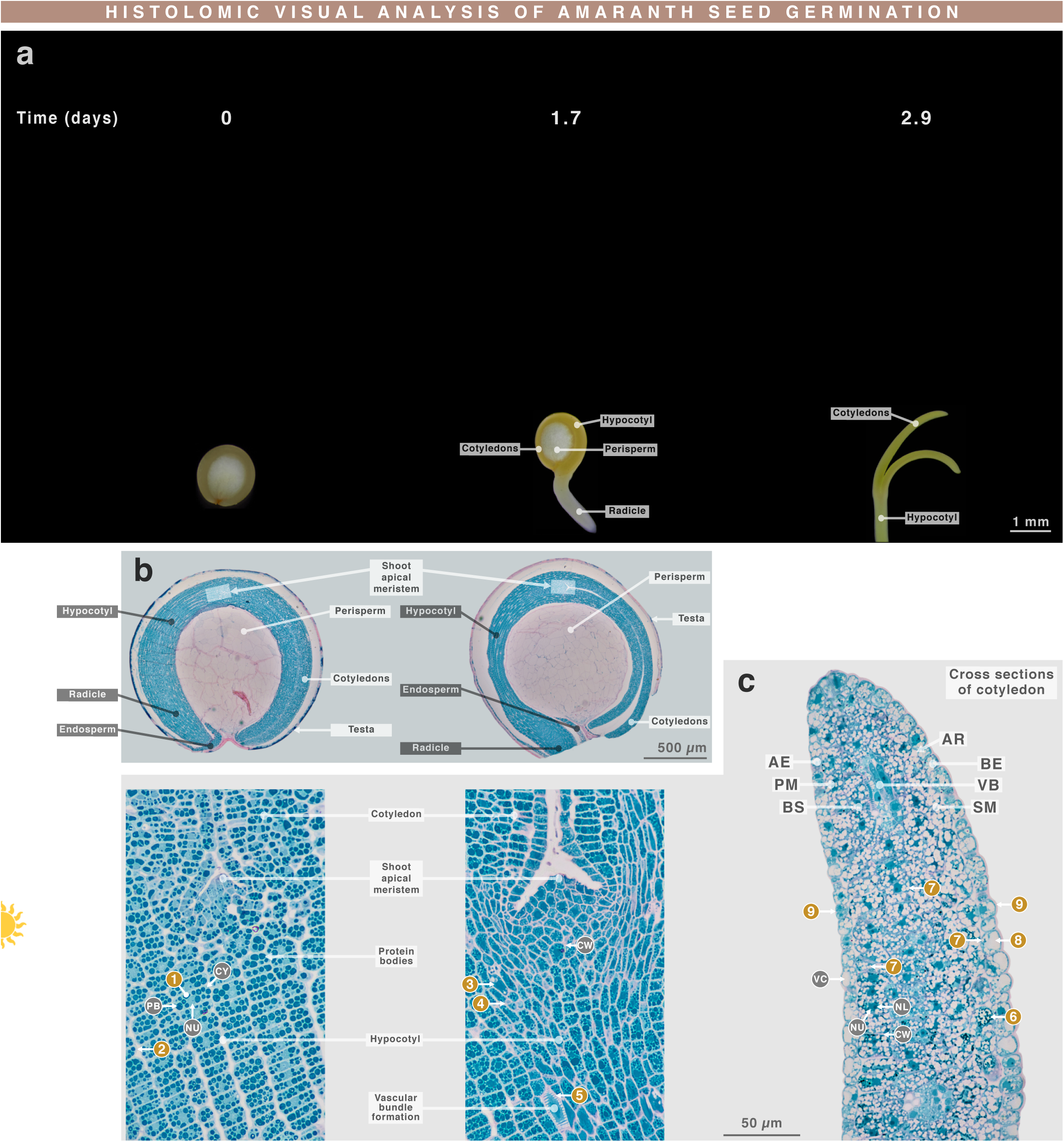
Histolomic visual analysis of *A. cruentus* seed germination. **a**, Polarization micrographs of *A. cruentus* seeds at different germination stages; days after sowing are indicated. **b**, Light micrographs of a whole seed (0 days) and germinating seed (1.7 days). **c**, Magnification of the boxed region in **b** and a cross section of the seedling cotyledon. Sections were thrichrome stained. Adaxial epidermis (AE), abaxial epidermis (BE), palisade mesophyll (PM), spongy mesophyll (SM), bundle sheath (BS), vascular bundles (VB), air spaces (AR). Cell wall (CW), vacuole (VC), chloroplast (CL), starch granule (ST), nucleus (NU), nucleolus (NL), cytosol (CY), mitochondrion (MI). The resin-based microtechnique was effective for amaranth seeds and seedlings. The embryo (cotyledons, hypocotyl, and radicle) stained turquoise, the testa stained magenta, and the perisperm remained unstained. Embryonic fusiform cells contained abundant protein bodies, nucleus, and cytosol (**1**), and their weakly stained CWs made cell boundaries difficult to distinguish (**2**). After germination, fusiform cells began degrading their protein bodies (**3**), thickened their CWs (**4**), and differentiated into conducting tissues such as tracheids (**5**). By three days, the cotyledon displayed all major leaf tissues—AE, PM, BS, VB, SM, BE, and some AR. Protein bodies had disappeared, although smaller, denser ones appeared (**6**). Starch granules accumulated in nearly all tissues (**7**), epidermal cells formed vacuoles (**8**), and BE developed a thicker outer wall than AE (**9**).

**SUPPLEMENTARY FIGURE 26.**
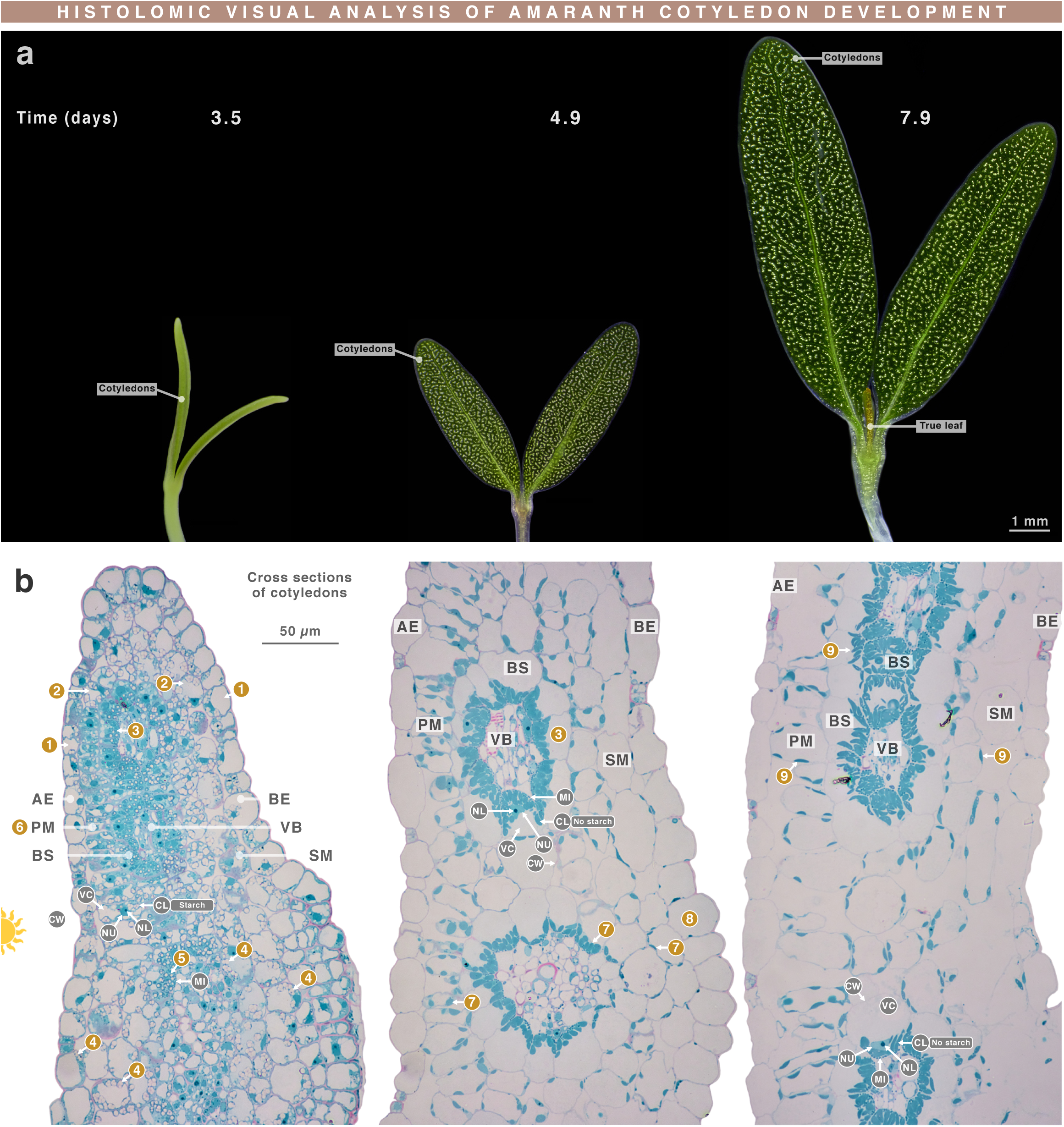
Histolomic visual analysis of *A. cruentus* cotyledon development. **a**, Polarization micrographs of *A. cruentus* cotyledons at different developmental stages; days after sowing are indicated. **b**, Light micrographs of cross sections from the cotyledons in **a**. Sections were thrichrome stained. Adaxial epidermis (AE), abaxial epidermis (BE), palisade mesophyll (PM), spongy mesophyll (SM), bundle sheath (BS), vascular bundles (VB), air spaces (AR). Cell wall (CW), vacuole (VC), chloroplast (CL), starch granule (ST), nucleus (NU), nucleolus (NL), cytosol (CY), mitochondrion (MI). By 3.5 days, all cells showed clear organelle development. Epidermal cells produced vacuoles first (**1**), followed by SM and PM cells (**2**), and finally BS cells (**3**). All tissues except VB accumulated starch granules **(4**). BS chloroplasts began adopting a centripetal arrangement (**5**), and some PM cells acquired their characteristic rod-like morphology (**6**). By 4.9 days, the cotyledon had a mature, leaf-like anatomy; all ST had disappeared (**7**) and epidermal cells had lost nearly all proteoplasm (**8**). By 7.9 days, true-leaf formation had begun, the cotyledon entered senescence, and chloroplasts adopted a slender, curved shape (**9**).

**SUPPLEMENTARY FIGURE 27.**
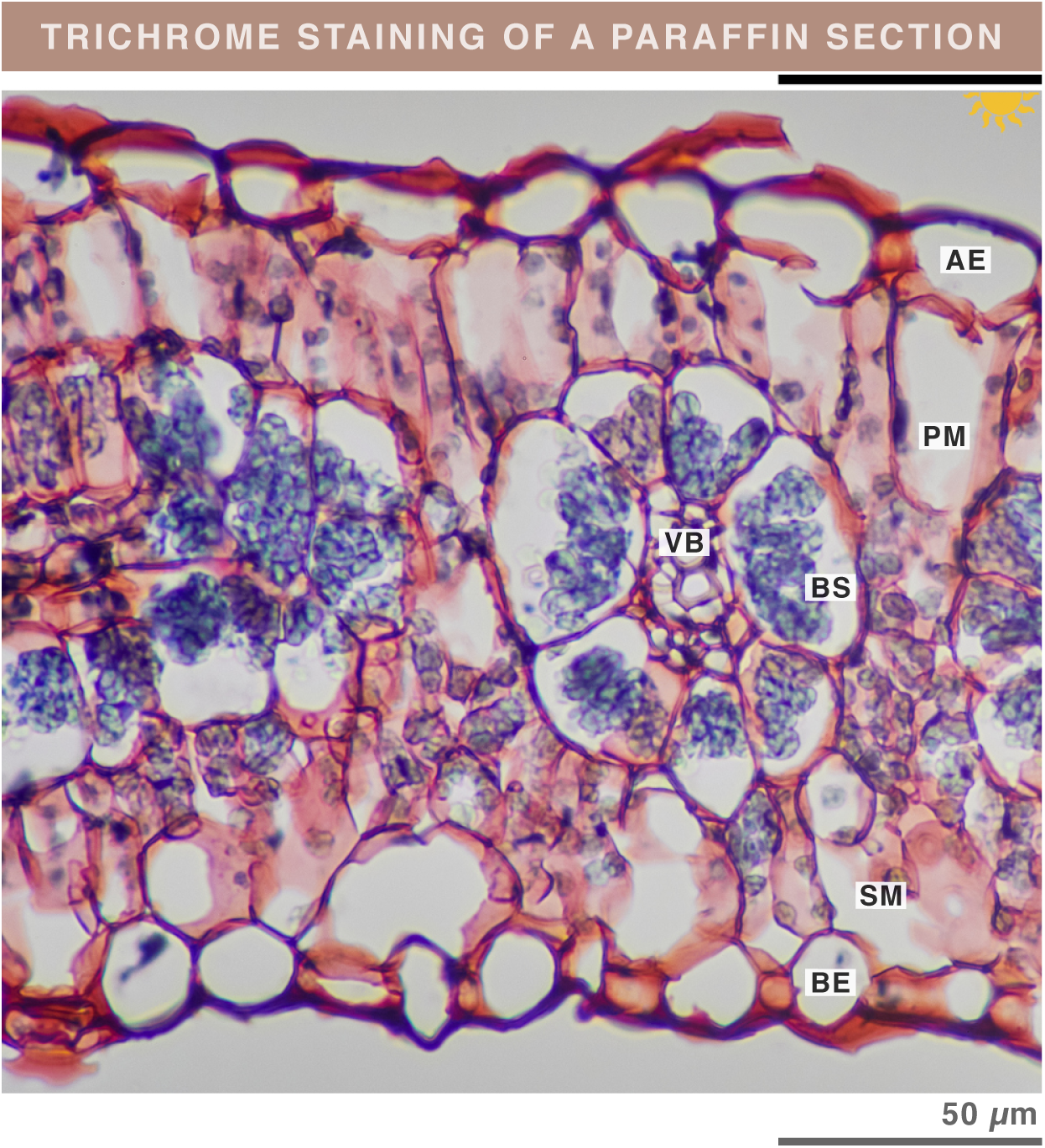
Trichrome staining also works on paraffin- embedded sections. Light micrograph of a paraffin-embedded *A. cruentus* leaf section stained with trichrome staining. The section was not deparaffinized. Trichrome staining was compatible with paraffin embedding. This section was 10 µm thick—20× thicker than resin sections—resulting in more intense staining. The strong coloration of starch granules partially masked the green of chloroplasts. In addition, PM and SM cells contained substantial cell-wall extensions, causing these regions to appear predominantly reddish.

